# Molecular responsiveness to testosterone is uncoupled from skeletal muscle mass and strength in males aged 18 - 80 years

**DOI:** 10.64898/2026.10.11.758372

**Authors:** Ross M. Williams, Megan Soria, Annabel Critchlow, Karel Van Belleghem, Viktor Engman, Andrew Garnham, Danielle Hiam, Glenn D. Wadley, Séverine Lamon

## Abstract

**Objective:** Testosterone exerts wide-ranging physiological effects, including the regulation of anabolic processes in skeletal muscle. However, its role in skeletal muscle ageing remains unclear. This study aimed to establish the association between endogenous testosterone levels and skeletal muscle phenotype across the adult lifespan and determine whether ageing modulates this relationship.

**Methods:** Eighty-one healthy males aged 18-80 underwent comprehensive phenotypic assessment. Body composition and muscle morphology were assessed using dual-energy X-ray absorptiometry (DEXA) and peripheral quantitative computed tomography (pQCT). Muscle strength was estimated leg press one-repetition maximum (e1RM). *Vastus lateralis* biopsies were used to assess fibre type, sequence the transcriptome and establish primary myocytes. Cell lines from young (n=5) and older (n=7) donors were cultured and compared under control and testosterone-treated conditions for cellular phenotype.

**Results:** Ageing was associated with lower free testosterone, muscle mass and strength, altered type IIa fibre characteristics, and widespread transcriptional changes in skeletal muscle. Free, but not total, testosterone was strongly associated with the muscle transcriptome, although neither fraction was associated with muscle mass or strength. In primary myocytes, testosterone increased androgen receptor abundance and nuclear localisation and enriched pathways involved in protein translation.

**Conclusion:** Testosterone is biologically active in male skeletal muscle, with its free fraction strongly associated with the transcriptome; however, these molecular effects do not translate into differences in muscle mass or strength. Testosterone may therefore play a primary role in establishing the male muscle phenotype during puberty, before transitioning to a more ancillary role in adult skeletal muscle.

## 1. Introduction

Skeletal muscle undergoes substantial remodelling over the course of the lifespan. At the functional level, ageing is accompanied by a loss of muscle mass, strength and contractile capacity (1). At the structural and cellular levels, ageing muscle undergoes significant changes in composition, fibre morphology and regenerative capacity (2-4). These changes do not occur concurrently or follow uniform linear trajectories (5). For instance, the rate at which males lose muscle mass accelerates greatly from the age of 60 onwards, and decrements in muscle strength typically precede those in muscle mass (6). It follows that conducting dichotomised comparisons between groups of young and older adult may overlook key periods of attenuation, stability or acceleration in muscle health across the adult lifespan.

Skeletal muscle does not age in isolation, and ageing is accompanied by substantial changes in the surrounding physiological environment, including the endocrine system (7). Of particular relevance to skeletal muscle is the age-related decline in circulating testosterone availability (8). Testosterone is the most abundant bioactive androgen and exerts wide-ranging physiological effects, including the regulation of anabolic processes in skeletal muscle (9). Most circulating testosterone is bound to sex hormone-binding globulin (SHBG) or albumin, and only a small fraction circulates in an unbound form (10). While SHBG-bound testosterone has restricted tissue availability, free testosterone, together with the more dissociable albumin-bound fraction, can more readily access tissue. However, total and free testosterone concentrations are often conflated despite providing different information regarding androgen exposure.

Despite the well-established anabolic effects of exogenous testosterone administration (11), the relationship between endogenous androgen concentrations and the skeletal muscle phenotype remains unclear. Associations between circulating testosterone and muscle mass, strength and related outcomes have been inconsistent, with interpretation complicated by methodological heterogeneity and potential confounding variables (12-15). Moreover, existing studies have predominantly examined total testosterone using linear models or comparisons between discrete age groups (16) and potential non-linear and age-dependent associations have not been comprehensively characterised. This is particularly important given that both skeletal muscle phenotypes and the endocrine environment change dynamically across adulthood, raising the possibility that androgen–muscle relationships themselves vary with age. Relationships between free testosterone and skeletal muscle outcomes remain comparatively underexplored.

The molecular mechanisms underlying these relationships are similarly poorly understood. Circulating testosterone concentrations represent only one component of androgen action and may not adequately reflect skeletal muscle responsiveness. The canonical actions of testosterone are mediated through its binding to the androgen receptor (AR), which, while, at rest, is maintained in a multiprotein complex within the cytoplasm (17). Testosterone-induced AR signalling regulates skeletal muscle growth and function *in vitro* (18-22) and *in vivo* (11, 23-26). Work from our group suggest that testosterone-induced myotube growth is primarily driven by canonical AR signalling rather than non-genomic pathways (18). Although older males retain whole-body anabolic responsiveness to exogenous testosterone (27), whether ageing alters the intrinsic molecular response of skeletal muscle cells to testosterone remains unclear.

This study aimed to comprehensively characterise circulating androgens, the skeletal muscle phenotype and the muscle transcriptome within a single, well-powered cohort spanning the male adult lifespan from 18-80, with rigorous control of relevant confounding variables. Non-linear models and age-interaction analyses were used to determine whether relationships between circulating androgens, muscle phenotype and gene expression varied across adulthood. To complement these *in vivo* analyses, which capture the influence of the systemic environment but cannot isolate direct cellular effects, we examined the molecular and morphological responses to testosterone in primary human muscle cells derived from young and older donors. This complementary approach allowed us to investigate whether ageing alters the intrinsic responsiveness of skeletal muscle cells to testosterone under controlled conditions.

## 2. Methods

### 2.1. Participants

Ethical approval was granted by the Deakin University Human Research Ethics Committee (2021-307). Eighty-one males aged 18–80 years, stratified by decade, were recruited from the local community via social media, posters and word of mouth. Eligible participants were biologically male, aged 18–80 years, with a body mass index (BMI) of 18–35 kg·m⁻². Exclusion criteria were type 2 diabetes; current cancer or <5 years remission; infectious bloodborne disease; anaesthetic allergies; smoking; use of dietary supplements likely to affect muscle synthesis; intellectual or mental impairment limiting understanding of the invasive procedures or capacity to provide informed consent; and a history of hormonal treatment or gender-affirming surgery. Before participation, all participants received a plain language statement detailing study procedures and provided written informed consent

### 2.2. Clinical and physiological measures

The outline of the study is depicted in **Supplementary Figure 1**. Prior to visit 1, participants abstained from alcohol, caffeine and vigorous exercise for a 48-hour period. At visit 1, leg strength was assessed using a 5-repetition-maximum (5RM) test using a leg press. Participants completed 2 warm-up sets of 8 repetitions, before completing up to 4 progressively heavier tests of 5 repetitions to establish their 5RM. Following each test, participants were asked to rate their exertion on a scale of 6-20 using the Borg Scale (28). The participants 5RM was recorded, and an estimated 1-repetition-maximum (e1RM) calculated using the following equation (29):

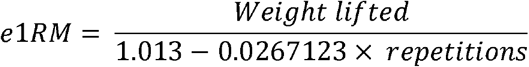

The following measures of body composition were assessed using dual energy x-ray absorptiometry (DXA; GE Lunar, Madison, WI): total body fat mass (TBFM), total body lean mass (TBLM), body fat percentage, appendicular lean mass (ALM), appendicular lean mass index (ALMI; calculated as ALM divided by height), bone mineral content, z-score. Peripheral quantitative computed tomography (pQCT; Stratec Medizintechnik GmBH, Pforzheim, Germany) scans of the upper thigh were used to assess muscle cross-sectional area (CSA; cm^2^), intramuscular fat area (cm^2^) and subcutaneous fat area (cm^2^).

At visit 2, no later than one month after the first visit, participants attended the laboratory in a fasted state, following a 48-hour period abstaining from alcohol, caffeine and vigorous exercise. A 5 ml blood sample was taken from the antecubital vein using a K2 EDTA blood collection tube (cat no. 32007-BD, McFarlane Medical, Victoria, Australia), centrifuged at 2500rpm at 4°C to isolate the plasma and stored at -80°C for further analysis. Hormone analysis was conducted by Monash Health Pathology. Total testosterone (TT), dihydrotestosterone (DHT), luteinizing hormone (LH), follicle stimulating hormone (FSH) and oestradiol (E2) were measured using high-performance liquid-chromatography mass-spectrometry (HP-LC-MS) (Triple Quad 5500, Sciex, Framingham, MA, USA). Sex hormone binding globulin was measured using a sequential two-step immunoenzymatic sandwich assay (UniCel DxI 800 Access Immunoassay System, Beckman Coulter, NSW, Australia). Free testosterone was calculated using the Vermeulen equation (30). The free androgen index (FAI) was calculated using the following equation:

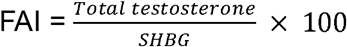

A muscle biopsy was then collected from the belly of the *vastus lateralis* muscle using the modified percutaneous Bergström technique (31). The skin was sterilised and anaesthetised using 1% Xylocaine (McFarlane Medical, Victoria, Australia). An incision was made in the skin and fascia using a scalpel blade. The muscle sample was removed, pre-cooled in liquid isopentane, and snap frozen in liquid nitrogen before long-term storage at -80°C. Approximately 100 mg of unfrozen tissue from each sample was immediately suspended in serum free Ham’s F10 nutrient mixture (cat. No. 11550-043, Thermo Fisher Scientific), ahead of isolating primary myocytes.

### 2.3. Control for confounding variables

Participants completed 7 days of diet and physical activity tracking in the interim period between visit 1 and visit 2. Participants recorded all food intake using the Easy Diet Diary app (Xyris Software, Brisbane, Australia), which was used to calculate average protein intake (g/kg body weight/day). Physical activity (minutes of moderate-vigorous physical activity; MVPA) was recorded using an ActiCal accelerometer wristwatch (Respironics Inc, Murraysville, PA). In addition, participants provided a detailed, self-reported written description of a typical week of physical activity, including type, duration, frequency and intensity of each exercise, which was used to categorise participants training status (**Supplementary Table 1**). Each reported activity was categorised by exercise modality using the Compendium of Physical Activities (32). Thresholds for ‘sedentary’, ‘recreational’ and ‘trained’ status for each modality were devised using previously published literature (33-35). Hybrid training status was assigned if criteria for more than one category were met, or if no single training modality contributed ≥ 50% of total weekly training minutes.

### 2.4. Skeletal muscle analysis

#### 2.4.1. Immunohistochemical staining and imaging

A subset of 43 participants was selected for skeletal muscle fibre-typing based on tissue availability and representing a minimum of n ≥ 5 per decade of age **(Supplementary Table 2)**. The sample size was determined via power calculation using the *pwr.f2.test* function in the *pwr* R package (36), with simulations based on Cuthbertson, Babraj (37), Léger, Derave (38) and Khanal, He (39).

*Vastus lateralis* muscle samples were immunohistochemically stained, processed and imaged to quantify muscle fibre type and size as described in (18) and described in Supplementary Methods.

#### 2.4.2. RNA Sequencing and Analysis

A subset of 65 participants was selected via stratified randomisation with random ordering (n = 7–15 per decade of age) **(Supplementary Table 3)**. RNA was extracted from approximately 10 mg of *vastus lateralis* tissue using an AllPrep DNA/RNA/miRNA Universal Kit (Cat No. 80224, Qiagen, Hilden, Germany). RNA quality and quantity were assessed using a TapeStation system (Agilent Technologies), with an RNA integrity number (RIN) of > 7 being considered acceptable for RNA-sequencing (Mean ± SD RIN: 8.35 ± 0.26). RNA-seq libraries were prepared from 1000 ng of RNA using the Ribo-Zero Gold protocol and Illumina TruSeq Stranded Total RNA protocol (Illumina, San Diego, CA). Samples were sequenced using the Illumina NovaSeq 6000 (Macrogen Oceania, NSW, Australia) with 150-bp paired-end reads and a sequencing depth of 25 million read pairs per sample. Reads were pseudo-aligned to the Ensembl human reference transcriptome (Ensembl 115 GRCh38.p14) (40) and transcript abundance quantified using Kallisto (v0.48.0) (41). Read quality was assessed using FastQC (v0.11.9, Babraham Institute, Cambridge, UK).

Transcriptomics analysis, clustering analysis and functional enrichment analysis were performed as described in Supplementary Methods.

Age x hormone interaction analyses were tested as described in Supplementary Methods.

#### 2.4.3. Isolation and culture of primary myoblasts

Myocytes were isolated from a sub-cohort of participants, with donor characteristics specified in **Supplementary Table 4**. Approximately 100 mg of muscle tissue was acquired from the *vastus lateralis* as described above, manually minced and suspended in serum-free Ham’s F10 nutrient mixture (cat. No. 11550-043, Thermo Fisher Scientific) (18, 42). The tissue was then dissociated three times in 0.05% Trypsin/EDTA (cat. no. 25300062, Thermo Fisher Scientific) at 35°C and cells suspended in proliferation media containing Ham’s F10 nutrient mixture, 20% foetal bovine serum (cat. No. A5670401, Thermo Fisher Scientific), 1% penicillin-streptomycin (cat. No. 15140122, Thermo Fisher Scientific), 0.05% amphotericin B (cat. no. 15290018, Thermo Fisher Scientific) and 25 ng ml^-1^ fibroblast growth factor (fGFb; cat. no. PHG0026, Thermo Fisher Scientific). The cells were then plated on to flasks pre-coated with extracellular matrix gel (cat. no. E1270, Sigma-Aldrich) and incubated at 37°C and 5% CO_2_ with proliferation media changed every 48 hours.

To isolate myocytes from other cell types, Magnetic Activated Cell Sorting (MACS) was performed as described in Supplementary Methods.

Purified myoblasts were seeded onto pre-ECM-coated six-well plates in proliferation media and left for 24 hours to adhere. Cell lines were grouped into young (18-30 years) and older (60-80 years) adult donor age groups. Proliferation based experiments were performed using 13 distinct cell lines (n = 6 young donors, n = 7 older adult donors). Twenty-four hours after seeding onto six-well plates, cells were treated with 100 nM testosterone (cat. No. 46923, Sigma-Aldrich) dissolved in ethanol (TESTO) or the equivalent volume of the vehicle control (CON).

Cells were treated with EdU at the onset of treatment for 24 hours to measure proliferation index. Cells were fixed for immunohistochemical staining 24 hours post-treatment in 4% PFA/PBS (cat. No. C005, ProSciTech) for 10 minutes. RNA was extracted 24 hours post-treatment and sequenced as described above.

Transcriptomics analysis, and functional enrichment analysis were performed as described in Supplementary Methods. Protein was extracted 48 hours post-treatment using 1x RIPA lysis buffer (cat. No. 20-188, Sigma-Aldrich) with 1% phosphatase and protease inhibitor cocktail (cat. no. 78440, Thermo Fisher Scientific). An outline of all primary myoblast-based experiments is presented in **Supplementary Figure 7.**

#### 2.4.4. Western Blotting

AR protein abundance was quantified by Western blotting, as detailed in the Supplementary Methods.

#### 2.4.5. Proliferation index, nuclei size and eccentricity

Myoblast proliferation, nuclear area and nuclear eccentricity were quantified using EdU incorporation and automated image analysis, as detailed in the Supplementary Methods.

#### 2.4.6. AR immunohistochemical staining and imaging

Whole-cell AR abundance and nuclear localisation were assessed by immunofluorescence microscopy and image analysis, as detailed in the Supplementary Methods.

#### 2.4.7. Telomere Length

Relative telomere length was determined by quantitative PCR using the telomere-to-single-copy-gene ratio, as detailed in the Supplementary Methods.

### 2.5. Statistical analysis

All statistical analyses were performed in RStudio (Version 2025.09.2+418) and R (Version 4.5.2). The distribution of muscle outcome and hormone data were visually inspected using histograms. Linear regression models were used to determine associations between age and circulating sex hormones, and age and muscle outcomes. Multiple linear regression models were used to determine associations between circulating sex hormones and AR expression and muscle outcomes, following adjustment for age, protein intake and training status where relevant. These covariates were selected as they have previously been shown to significantly affect muscle outcomes (43-46). Participants missing hormone or muscle outcome data were excluded from the relevant models. Missing protein intake data were imputed using the R *MissForest* package (n = 15; 5.4% of the cohort) (47, 48). Linear, polynomial (2^nd^) and generalised additive models were compared for best fit using Akaike Information Criterion (AIC), with the model with the lowest AIC being the preferred fit. Heteroscedasticity and non-linearity were assessed via visual inspection of residual vs fitted plots, and Shapiro-Wilk and Breusch-Pagan tests. Normality was visually assessed using quantile-quantile plots. Multicollinearity was assessed using variance inflation factors (VIF), with all models reporting VIF < 1.4. Linear models were used in all cases but the fibre-type proportion outcomes, which were modelled using beta regressions using the *betareg* R package. Significance was set at *p* < 0.05.

To explore whether associations between circulating sex steroid hormones and muscle outcomes varied with age, interaction effects between each hormone and age were added to the linear and beta regression models. Given the exploratory nature of this analysis, p-values were not adjusted for multiple comparisons. Regression coefficients from both linear and beta regression models were standardised to enable comparison of effect sizes across hormones and outcomes. Standardised coefficients were visualised using the R *ggplot2* package (49) via bar plots and heatmaps using a fixed colour scale ranging from -1 to 1, with values outside this range truncated to facilitate comparisons. Standardised coefficients were used for visualisation only, with all statistical analyses and inferences based on unstandardised coefficients.

Statistical analyses related to cell experiments are described in Supplementary Methods.

## 3. Results

### 3.1. Participant characteristics

Eighty-one males, representing every decade of age between 18-80, completed the study. Participant characteristics are detailed in **Table 1**. The subsets of participants who underwent muscle fibre histological analysis and RNA-sequencing, respectively, are characterised in **Supplementary Table 2 and 3.**

**Table 1.** Participant characteristics. BMI: Body mass index; ALM: Appendicular lean mass; ALMI: Appendicular lean mass index; CSA: Cross-sectional-area; 1RM: One-repetition-maximum; e1RM: Estimated one-repetition maximum; MVPA: Moderate-vigorous physical activity. Data is mean ± SD

| Variable | Overall | Aged 18-29 | Aged 30-39 | Aged 40-49 | Aged 50-59 | Aged 60-69 | Aged 70-79 |
| --- | --- | --- | --- | --- | --- | --- | --- |
|  | N = 81 | N = 15 | N = 13 | N = 13 | N = 10 | N = 18 | N = 12 |
| **Body composition** |  |  |  |  |  |  |  |
| Weight (kg) | 79.8 (11.8) | 75.4 (9.4) | 80 (13.5) | 88 (14.4) | 82.4 (7.6) | 80.8 (11) | 72.6 (8) |
| Height (m) | 176.8 (8) | 177.8 (6.5) | 176.7 (8.5) | 177.7 (10.5) | 181.2 (8.7) | 174.2 (7.1) | 174.6 (6.4) |
| BMI (kg/m<sup>2</sup>) | 25.5 (3.3) | 23.8 (2.6) | 25.6 (3.2) | 27.8 (3.7) | 25.3 (3.4) | 26.6 (3.1) | 23.8 (1.7) |
| Body fat (%) | 24.1 (7.8) | 16.7 (5.3) | 23.5 (6.3) | 28.4 (8.9) | 22.8 (7) | 28.3 (7) | 24.4 (5.4) |
| Total body lean mass (kg) | 58 (8.6) | 59.1 (6.2) | 58.7 (10.7) | 61.7 (12.1) | 61.7 (8) | 55.1 (6.7) | 53.1 (4.1) |
| Total body fat mass (kg) | 18.9 (8) | 12.6 (5.9) | 18 (5.5) | 24.5 (10.8) | 18.2 (6.9) | 22.1 (7) | 17.1 (5.3) |
| Appendicular lean mass (kg) | 26.9 (4.1) | 28.4 (3.1) | 28.2 (5.8) | 28.6 (3.9) | 28 (2.3) | 25.2 (3.6) | 23.4 (2.3) |
| ALM Index (kg/m<sup>2</sup>) | 8.6 (1) | 9 (0.8) | 9 (1.5) | 9 (0.7) | 8.5 (0.7) | 8.3 (1) | 7.7 (0.5) |
| Bone Mineral Content (kg) | 3.1 (0.4) | 3.1 (0.3) | 3.2 (0.6) | 3.2 (0.4) | 3.3 (0.4) | 3 (0.4) | 3.1 (0.4) |
| **Muscle size and composition** |  |  |  |  |  |  |  |
| Thigh muscle CSA (cm<sup>2</sup>) | 134.5 (24.4) | 148.9 (17.3) | 148.8 (33.7) | 141.6 (19.9) | 137.2 (11.9) | 122.1 (17.6) | 110 (13.9) |
| Intramuscular fat (cm<sup>2</sup>) | 4.4 (3) | 2.4 (2.2) | 4.1 (2.9) | 6.1 (4.1) | 4.4 (2) | 5.6 (3.1) | 3.9 (2) |
| Subcutaneous fat (cm<sup>2</sup>) | 35.8 (17.3) | 32.4 (22.8) | 40.3 (13.8) | 44 (21.4) | 29.5 (10) | 40 (14.4) | 25.4 (10.4) |
| **Muscle strength** |  |  |  |  |  |  |  |
| Leg press e1RM (AU) | 276.4 (79) | 334.7 (74.1) | 315.3 (90) | 291.2 (78) | 279.2 (52.5) | 237.8 (50) | 201 (37.8) |
| e1RM normalized to LLM (AU/kg) | 14 (3) | 16.3 (3) | 15.2 (2.8) | 13.8 (2.9) | 13.8 (2.7) | 12.8 (2.1) | 11.8 (2.8) |
| E1RM normalized to muscle CSA (AU.cm2) | 2 (0.3) | 2.2 (0.4) | 2.1 (0.3) | 2 (0.4) | 2 (0.3) | 1.9 (0.2) | 1.8 (0.3) |
| **Circulating sex hormones** |  |  |  |  |  |  |  |
| Total testosterone (nmol/L) | 21.6 (7.3) | 24.1 (10.1) | 23.4 (7.2) | 21.2 (7.5) | 18.6 (4.6) | 18.7 (4.7) | 23.8 (7.3) |
| Free testosterone (pmol/L) | 412.8 (144.1) | 500 (189.8) | 509 (147.1) | 430.9 (103.8) | 356.8 (121.2) | 330.7 (78) | 351.4 (88.2) |
| Free androgen index (%) | 57.3 (24.1) | 69.3 (28.5) | 71.7 (20.4) | 63.1 (18) | 55.6 (33.7) | 44.6 (13.8) | 41.3 (10.8) |
| Dihydrotestosterone (nmol/L) | 1.7 (0.7) | 1.8 (0.7) | 1.8 (0.6) | 1.6 (0.8) | 1.3 (0.5) | 1.5 (0.6) | 2.1 (0.9) |
| Sex hormone binding globulin (nmol/L) | 42.8 (19.3) | 37.4 (17) | 34.9 (12.6) | 37.5 (20.8) | 42.2 (22.1) | 45.1 (15.1) | 60.5 (21.4) |
| Estradiol (pmol/L) | 114.8 (29.5) | 124.6 (31.3) | 107 (33.5) | 108.3 (32.5) | 102.2 (17.2) | 125 (29) | 112.2 (24.1) |
| **Lifestyle factors** |  |  |  |  |  |  |  |
| MVPA (mins/day) | 41.9 (33.2) | 48.8 (37.8) | 46.9 (25) | 44.6 (43) | 45.6 (24.8) | 31 (33.8) | 37.8 (31.9) |
| Dietary protein intake (g/kg/day) | 1.4 (0.5) | 1.9 (0.7) | 1.5 (0.3) | 1.2 (0.3) | 1.3 (0.2) | 1.2 (0.5) | 1.3 (0.3) |
| Most prevalent training type | hybrid | hybrid | hybrid | sedentary | endurance | endurance | body weight exercise |

### 3.2. Effect of age on circulating androgens, muscle characteristics and the muscle transcriptome

Age was not associated with total testosterone (TT), dihydrotestosterone (DHT) or oestradiol (E2) concentrations. Free testosterone (FT) concentrations and the free androgen index (FAI) significantly decreased with age (both *p* < 0.001) (**Figure 1A**).

**FIGURE 1.**
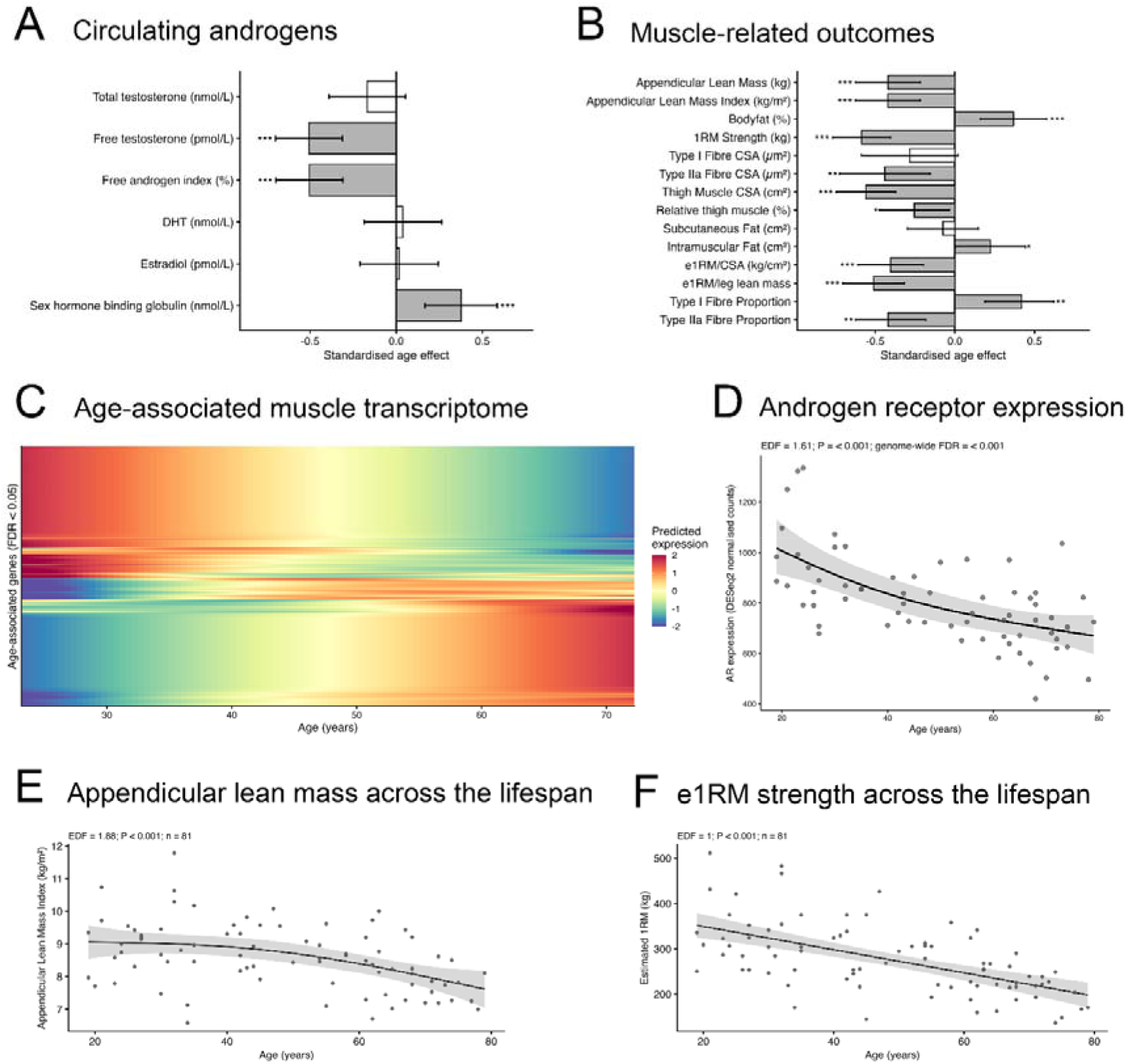
Effect of age on circulating androgens, muscle characteristics and the muscle transcriptome. Standardised associations between age and (A) circulating hormone concentrations and (B) muscle-related outcomes in healthy adult males. Bars represent standardised regression coefficients (β) and error bars indicate 95% confidence intervals. Filled bars denote statistically significant associations (P < 0.05). Asterisks denote level of statistical significance (*P < 0.05, **P < 0.01, ***P < 0.001). DHT: Dihydrotestosterone; e1RM: Estimated one-repetition-maximum; 1RM: One-repetition-maximum; CSA: Cross-sectional area. Heatmap of age-associated genes identified using gene-wise generalised additive models (FDR < 0.05). Each y-axis row represents a gene significantly associated with age (FDR < 0.05). X-axis represents gene expression across the lifespan. Colour intensity indicates scaled predicted gene expression, with warmer colours representing relatively higher expression and cooler colours representing relatively lower expression within each gene (C). Expression levels of AR mRNA modelled against the age continuum (D). ALMI modelled *against the age continuum (E). e1RM modelled against the age continuum (F)*.

Appendicular lean mass (ALM), ALM index (ALMI), thigh muscle cross-sectional area (CSA), thigh muscle percentage (%), absolute leg press estimated 1-repetition maximum (e1RM) and e1RM relative to thigh muscle CSA and leg lean mass significantly decreased with age (all *p* < 0.001). Intramuscular fat infiltration (*p* < 0.05), but not subcutaneous fat, was positively associated with age. Type IIa muscle fibre CSA and muscle fibre proportion were negatively associated with age (both *p* < 0.01). Type I muscle fibre proportion was positively associated with age (*p* < 0.01) (**Figure 1B**). Lean muscle mass and muscle strength, as measured by ALMI and e1RM, respectively, decreased by approximately 3% and 7% and per decade, respectively (**Figure 1E and 1F).**

In a subsample of 65 males stratified by decade of age (**Supplementary Table 3**), age was significantly associated (FDR < 0.05) with the expression of 2819 genes, 1764 of which were approximately linear (edf < 1.2) and 1055 of which were non-linear (**Figure 1C**). Of particular interest, age was negatively associated with AR gene expression levels (FDR < 0.01) (**Figure 1D**). REACTOME and Gene Ontology analysis identified that genes positively associated with age were enriched for biological pathways including cell adhesion, RNA splicing, mRNA processing and the cellular response to endogenous stimulus. Genes negatively associated with age were enriched for pathways including DNA methylation, gluconeogenesis and mitochondrial membrane organisation (**Supplementary Figure 3A-B**). Genes were grouped into eleven clusters based on their expression trajectories across the lifespan, and individual pathway analysis was subsequently conducted for each cluster (**Supplementary Figure 4**).

### 3.3. Associations between circulating androgens, muscle characteristics and the muscle transcriptome

Following adjustment for age, protein intake and training status, TT was negatively associated with body fat percentage and intramuscular fat (both *p* < 0.01) but not with any other measures of body composition or muscle strength. FT, FAI, DHT and E2 were not associated with any measures of body composition or muscle strength (**Figure 2A**). FT, FAI and E2 were positively associated with Type IIa muscle fibre proportion (all *p* < 0.05) and negatively associated with Type I muscle fibre proportion (FT: *p* < 0.01; FAI and E2: *p* < 0.05). FT and FAI were positively associated with Type I muscle fibre CSA (both *p* < 0.05) (**Figure 2B**).

**FIGURE 2.**
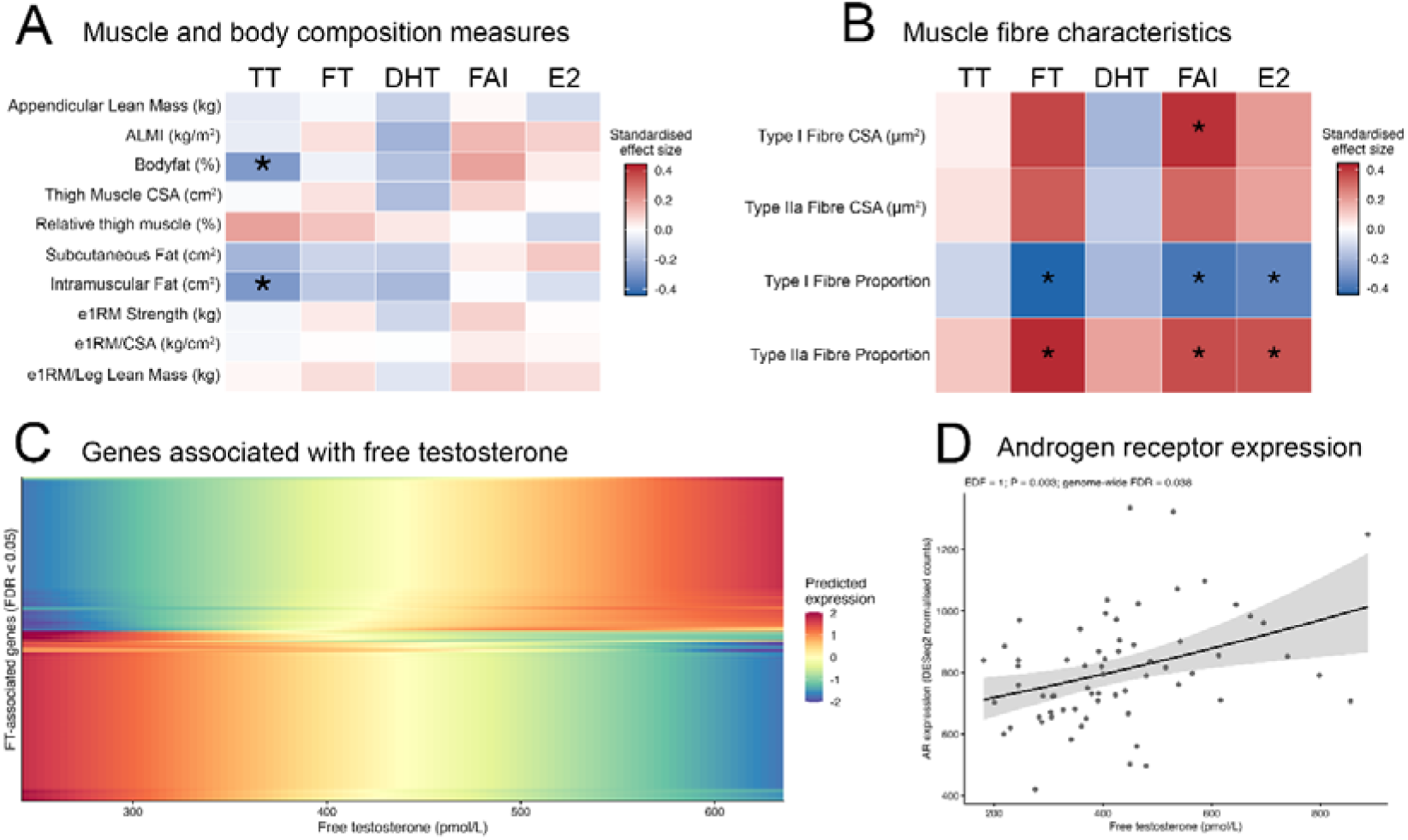
Associations between circulating androgens, muscle characteristics and the muscle transcriptome. Standardised associations (β) between circulating sex steroid hormone concentrations and (A) measures of body composition, muscle size, muscle strength and (B) muscle fibre characteristics adjusted for age, protein intake and training status. Cell colour represents the magnitude and direction of the standardised effect size. Asterisks denote statistically significant associations (p < 0.05). DHT: Dihydrotestosterone; E2: Estradiol; FAI: Free androgen index; FT: Free testosterone; TT: Total testosterone; e1RM: Estimate one-repetition-maximum; 1RM: One-repetition-maximum; CSA: Cross-sectional area. Heatmaps of testosterone-associated genes identified using gene-wise generalised additive models (FDR < 0.05). Heatmap shows genes significantly associated with free testosterone. Each row represents a significantly associated gene, and columns represent free testosterone concentrations across the modelled hormone continuum. Colour intensity indicates scaled predicted gene expression, with warmer colours representing relatively higher expression and cooler colours representing relatively lower expression within each gene (C). Expression levels of AR mRNA modelled against the free testosterone level continuum (D).

Total testosterone was not associated with the expression of any gene. In contrast, free testosterone was associated with the expression of 1509 genes, 1131 of which were approximately linear (edf < 1.2) and 378 of which were non-linear (**Figure 2C).** Of particular interest, free testosterone was positively associated with AR gene expression levels (FDR < 0.01) (**Figure 2D**). Genes positively associated with free testosterone concentrations were enriched for various glycolytic and glucose metabolic processes. Genes negatively associated with free testosterone were enriched for circulatory system development, negative regulation of viral genome replication and the response to peptides (**Supplementary Figure 5A-B**). Genes were grouped into four clusters based on their expression trajectories relative to free testosterone concentrations (**Supplementary Figure 6**), and individual pathway analysis was subsequently conducted for each cluster.

There was no significant age x total testosterone, or age x free testosterone interaction effects for any genes.

### 3.4. Effect of donor age and testosterone treatment on human primary myoblast phenotype

We next investigated the effect of testosterone treatment on phenotypical aspects of human primary myoblasts. We first sought to determine whether donor age had any significant effect on myoblast characteristics. Experimental timelines are presented in **Supplementary Figure 7**. There was no significant effect of donor age on nuclear area, nuclear eccentricity, proliferation index following a 4-hour or 24-hour EdU pulse, whole myoblast AR intensity, AR protein content, nuclear AR localisation or myoblast telomere length (**Supplementary Figure 8A-J, Supplementary Figure 9**).

Testosterone treatment did not alter myoblast nuclear area, nuclear eccentricity or proliferation index following a 4-hour or 24-hour EdU pulse (**Figure 3A-D**), but significantly increased whole myoblast AR intensity (determined by immunohistochemistry) (**Figure 3E**), AR nuclear localisation (determined by immunohistochemistry) (**Figure 3F-G**) and AR protein content (determined by Western Blot) (**Figure 3H-I**) in cell lines from both young and old donors (all *p* < 0.001).

**FIGURE 3.**
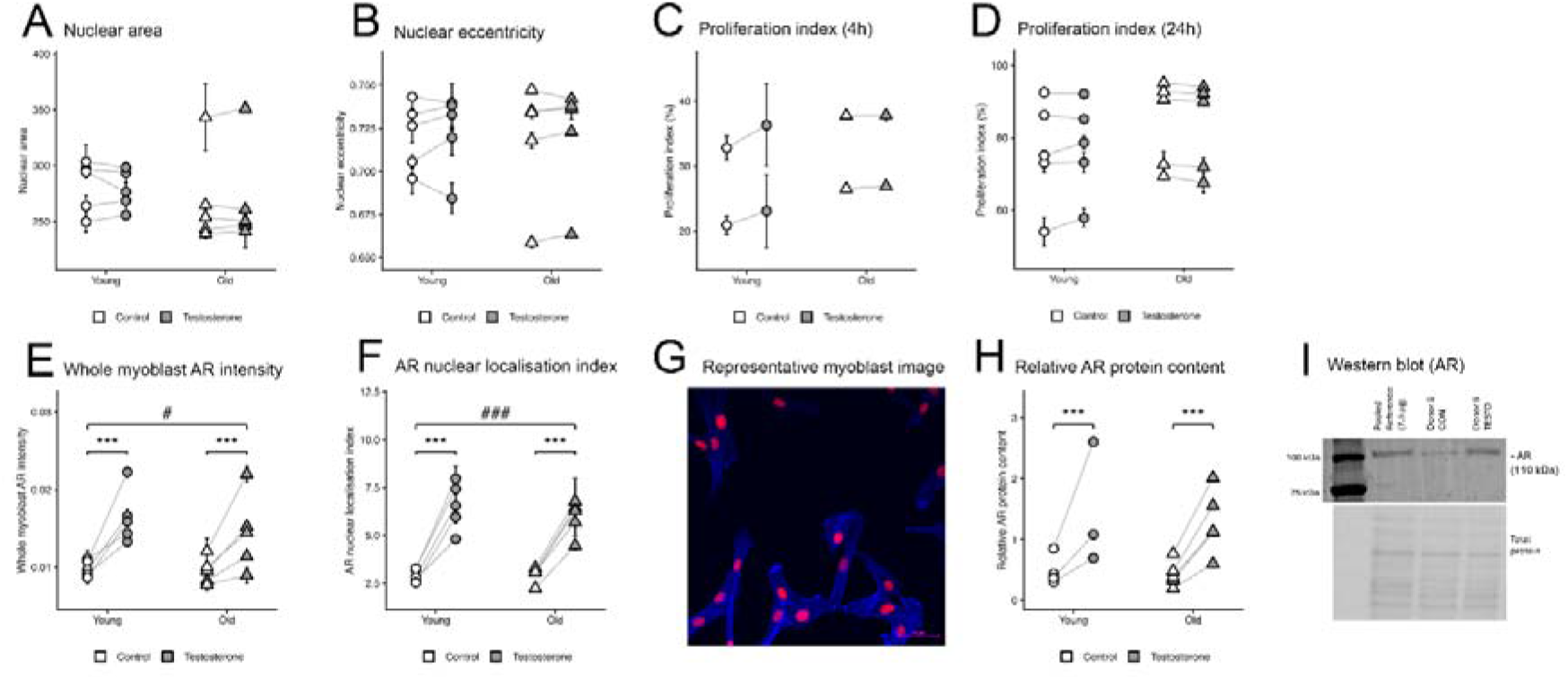
Effects of testosterone treatment on primary human myoblast characteristics in young and older donors. Effects of testosterone (TESTO) or vehicle control (CON) treatment on (A) nuclear area and (B) nuclear eccentricity in primary human myoblasts derived from young (n = 5) and older (n = 5) donors; (C) proliferation index following a 4-h EdU pulse (n = 2 per age group); (D) proliferation index following a 24-h EdU pulse (n = 5 per age group); (E) whole-myoblast androgen receptor (AR) intensity and (F) AR nuclear localisation index derived from young (n = 5) and older (n = 5) donors. (G) Representative image of myoblasts stained for phalloidin (blue), DAPI (red) and total AR protein (green) following 24 hours TESTO treatment. Scale bar = 50 µm. (H) Relative AR protein content in myoblasts derived from young (n = 3) and older (n = 5) donors following 48 hours CON or TESTO treatment. (I) Representative AR western blot membrane following 24 hours CON or TESTO treatment. AR bands are shown at approximately 110 KdA. Data are presented as individual donor-level means, error bars indicate standard deviation (SD) between replicate wells, connecting lines indicate donor treatment response. Error bars indicate between-well standard deviation. *** indicates a significant treatment effect (p < 0.001). # and ### indicate significant age × treatment interaction effects at p < 0.05 and p < 0.001, respectively.

In addition, there was a significant treatment x age interaction effect for whole myoblast AR intensity (*p* < 0.05) (**Figure 3E**) and AR nuclear localisation (*p* < 0.05) (**Figure 3F**), indicating testosterone effect was stronger in myoblasts from young than in myoblasts from old donors.

Finally, we examined the effects of donor age and testosterone treatment on the myoblast transcriptome. Age was associated with the differential expression of 11 transcripts (FDR <0.05, 0.06% of all detected transcripts; **Supplementary Figure 10**), and no significant age × testosterone treatment interactions were identified for any transcripts (**Supplementary Figure 11**). Therefore, cell lines were pooled across age groups for subsequent analyses.

Testosterone treatment significantly altered the expression of only five transcripts (FDR <0.05, 0.025% of all detected transcripts; **Figure 4A**), including *MYF5*, a myogenic regulatory factor previously shown to be responsive to androgen treatment in human skeletal muscle cells (50). Reactome enrichment analyses identified an overrepresentation of pathways related to translation, translation initiation and termination, and ribosomal processes among transcripts upregulated by testosterone (**Figure 4B**).

**FIGURE 4.**
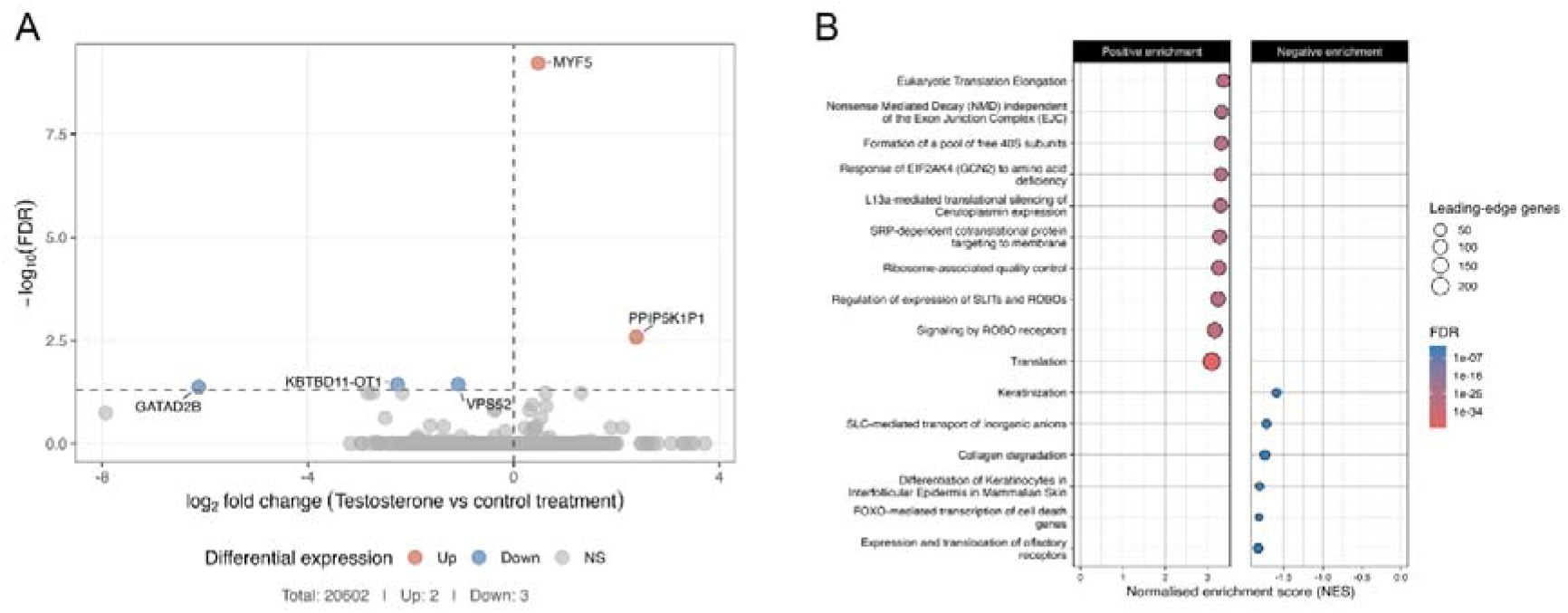
Effects of testosterone treatment on the primary human myoblast transcriptome. Volcano plot of differentially expressed transcripts in primary human myoblasts (N = 10) following 24hrs 100nM testosterone treatment. The x-axis represents log fold change (testosterone/control) and the y-axis represents −log adjusted P value. Significantly upregulated and downregulated transcripts (adjusted P < 0.05) are shown in red and blue, respectively. (B) Reactome pathway enrichment analysis of genes associated with testosterone treatment. Pathways with positive normalized enrichment scores (NES) are shown as activated and those with negative NES as suppressed. Circle size represents the number of genes contributing to each pathway and colour represents the adjusted P value.

## 4. Discussion

This study aimed to investigate the relationship between age, circulating sex steroid hormone concentrations and the skeletal muscle phenotype and transcriptome across the adult male lifespan, and to probe the potential origins of these relationships in a tissue culture model. Overall, age was negatively associated with free but not total testosterone concentrations, skeletal muscle mass, strength and the fast-twitch muscle phenotype. These relationships were accompanied by widespread age-associated transcriptomic variation. Circulating sex steroid hormone concentrations did not exhibit associations with skeletal muscle mass or strength following adjustment for relevant covariates, including age. However, total testosterone concentrations were associated with distinct aspects of skeletal muscle quality and composition, and free testosterone concentrations were strongly associated to the muscle transcriptome. Human primary myoblasts did not exhibit distinct characteristics of donor age under basal conditions; though donor age did influence the effect of testosterone treatment on AR expression and localisation.

Consistent with well-established characteristics of skeletal muscle ageing, we report negative associations between age and various measures of muscle mass and strength (2, 5, 51-56) along extensive variations in the molecular profile (57-60). A novel finding, however, was that age-related associations with the transcriptomic profile were not exclusively linear, with many genes exhibiting complex, non-linear expression trajectories across the male lifespan. Changes in circulating androgen concentrations across the lifespan are well recognised, although their magnitude, determinants and physiological significance remain debated. Large-scale epidemiological studies often report moderate declines in total testosterone and DHT with age, albeit with highly variable magnitudes of effect (7, 61-63). In contrast, our results indicate a negative association of age with free testosterone concentrations, a component of the bioavailable fraction of testosterone, and the FAI, but not total testosterone or DHT concentrations. Typically, studies reporting smaller, non-significant effect sizes are those implementing rigorous control for covariates such as obesity and other comorbidities (64, 65). As such, our findings are consistent with literature reflecting the effects of healthy biological ageing on the male hormonal profile once corrected for the relevant variables.

Neither total nor free testosterone were independently associated with any measure of muscle mass or strength following adjustment for age, physical activity and protein intake. Across the adult lifespan, the regulation of skeletal muscle is a multifactorial process. As such, the influence of testosterone concentrations within the normal range on adult skeletal muscle is likely much smaller when isolated from other factors (66, 67), especially age. In addition, the rise in circulating testosterone during male puberty is an important driver of skeletal muscle accrual and contributes to the acceleration of sex differences in muscle mass and strength during adolescence (68). However, the relationship between circulating testosterone and muscle mass appears to weaken toward completion of puberty (69), suggesting that endogenous testosterone may be more important for establishing the male muscular phenotype than for determining variation in muscle mass and strength following puberty. Our transcriptomic findings further support this interpretation, and total testosterone did not exhibit any significant relationship with the adult muscle transcriptome.

Therefore, the perceived associations between total endogenous testosterone concentrations and skeletal muscle mass and strength may be largely attributable to shared associations with age, physical activity and diet. Collectively, these findings suggest that androgen availability, rather than total content, may influence the molecular and contractile phenotype of skeletal muscle without being a major determinant of whole-muscle mass or maximal strength. A rate-limiting step in this process may be androgen receptor availability. We recently used a meta-analytic approach to confirm that skeletal muscle androgen receptor gene expression declines with age (70). Its positive association with the free testosterone fraction suggests a potential feedforward relationship between androgen availability and receptor expression.

An unexpected finding of this study was the relationship between free, but not total testosterone and the muscle transcriptome. This distinction is biologically plausible, as total testosterone includes the fraction bound to SHBG whereas free testosterone more closely reflects the fraction readily available to skeletal muscle. This may be particularly relevant across the male lifespan, as circulating SHBG concentrations increase with age, potentially uncoupling total testosterone concentrations from androgen bioavailability. Free testosterone may therefore provide a more physiologically relevant indicator of androgen exposure at the tissue level and, consequently, of androgen-sensitive transcriptional activity. In combination with our previous work (70), the association between free testosterone and AR mRNA expression also raise the hypothesis that androgen availability may influence muscle transcription through both ligand availability and modulation of receptor abundance. The > 1500 genes associated with free testosterone demonstrated strongest enrichment for pathways involved in substrate metabolism and immune defence. Given that transcriptomic profiling was performed in whole skeletal muscle tissue, the immune-related signature likely reflects the integrated transcriptional contribution of myofibres and non-myofibre cell populations, including resident and infiltrated immune cells (71). These findings may therefore indicate an association between circulating free testosterone and the local immune/inflammatory environment of skeletal muscle, rather than a direct androgen-mediated immune response within muscle fibres.

To probe whether some of the associations observed in humans may stem from an age-modulated response to testosterone, we next investigated the interplay between age and testosterone treatment on phenotypical aspects of human primary myoblasts. In line with our recent transcriptomics results in human myotubes (42), we report a lack of effect of donor age on various aspects of proliferating human myoblasts, including proliferation rate, telomere length, nuclear morphology, nuclear localisation or the transcriptome at baseline or following testosterone treatment. These findings are consistent with prior research reporting no difference in the proliferative capacity of myoblasts isolated from young and older adult donors (72-75); although donor age may attenuate proliferation over extended periods of cultivation (76). Other aspects of myogenic cell function, such as greater incidence of senescence, mitochondrial dysfunction and impaired differentiation have been observed in cells isolated from older adult donors (72, 73, 77, 78), but these observations were made in differentiating myotubes, rather than myoblasts.

Testosterone administration did not alter nuclear morphology or proliferative capacity in myoblasts. An accelerated cell-cycle progression is not a hallmark of testosterone’s action in skeletal muscle. Rather, myonuclear accretion is driven by increased satellite cell activation and progression through the myogenic program (24, 79-82). Testosterone administration however increased AR protein abundance and nuclear translocation. Similar effects have previously been reported by our group in young female myotubes (18) and *in vivo* in human skeletal muscle (83). These findings indicate that cells in the early proliferative stages of the myogenic lineage are responsive to testosterone and exhibit receptor localisation consistent with activation of the canonical AR signalling pathway described in mature myocytes (84-86).

While myoblasts isolated from older adults did not exhibit distinct characteristics of donor age under basal conditions, donor age did influence the response to testosterone treatment, with older donor cells exhibiting attenuated changes in whole myoblast AR intensity and AR nuclear localisation. Donor age did not modify the testosterone-induced increase in total AR abundance when measured via Western Blot, which might indicate that the observed donor-age dependent response extends beyond changes in total receptor abundance, and instead encompasses the cellular handling and localisation of the AR. Our study is not the first to suggest that cultured myoblasts may retain donor-specific phenotypic characteristics that become apparent in their response to physiological stimuli rather than under basal conditions. However, evidence remains limited and has largely focused on metabolic phenotypes, with features of donor insulin resistance previously shown to persist in both myoblasts (87) and myotubes (88).

At the transcriptomic level, the effects of testosterone treatment were limited but recapitulated earlier findings from a similar intervention in primary human myotubes assessed using a candidate-gene approach (50). In particular, MYF5 was the most strongly regulated transcript following testosterone treatment, consistent with previous observations that testosterone and DHT increase MYF5 expression in primary human myotubes without concomitant changes in MYOD, MYOG or AR expression (50). The absence of a broader transcriptional response aligns with evidence that several actions of testosterone in muscle are mediated through post-transcriptional and post-translational mechanisms, such as miRNAs (50). Nevertheless, despite the small number of individually differentially expressed transcripts, pathway analysis revealed coordinated enrichment of genes involved in translation initiation, elongation and termination, ribosomal function and co-translational protein targeting. This confirms that testosterone elicits a broader transcriptional programme associated with translational capacity, consistent with its established effects on muscle protein synthesis. Importantly, the absence of an age x testosterone interaction indicates that donor age did not detectably modify the transcriptional response to testosterone under these experimental conditions, further supporting the concept that age-related changes in muscle cells and androgen-mediated regulation represent at least partly independent processes.

This study highlighted extensive variation in the skeletal muscle phenotype and transcriptome across the adult male lifespan, encompassing skeletal muscle mass, strength, architecture and their associated regulatory pathways. However, despite these age-associated differences, circulating testosterone concentrations were not widely associated with skeletal muscle mass or strength; instead, we found that testosterone exhibited specific relationships with aspects of skeletal muscle quality and composition, and free testosterone was more closely related to skeletal muscle’s contractile architecture and the skeletal muscle transcriptome. These findings challenge the standing that physiological variation in endogenous testosterone concentrations is a major determinant of skeletal muscle mass and strength in adult males and highlights that covariates such as age and physical activity may incur confounded associations misattributed to testosterone concentrations. Our results indicate that, in adult males, testosterone concentrations may occupy a more specific role in the maintenance of the skeletal muscle architecture. Alternatively, testosterone may play a more central role during puberty, establishing the musculoskeletal phenotype, before transitioning to a more ancillary role in later life, modifying its structural and functional characteristics.

## Acknowledgments

We sincerely thank all participants for generously volunteering their time to take part in this study. R

## Authors’ contributions

Ross M. Williams: Investigation, Formal analysis, Writing – original draft, Writing – review & editing. Megan Soria: Investigation, Formal analysis, Writing – original draft, Writing – review & editing. Annabel Critchlow: Investigation, Writing – review & editing. Karel Van Belleghem: Investigation, Writing – review & editing. Viktor Engman: Investigation, Writing – review & editing. Andrew Garnham: Investigation, Writing – review & editing. Danielle Hiam: Investigation, Formal analysis, Writing – review & editing. Glenn D. Wadley: Investigation, Writing – review & editing. Séverine Lamon: Conceptualization, Funding acquisition, Investigation, Writing – original draft, Writing – review & editing.

## Data availability

The R code for analysis can be found at https://github.com/ROSWILLIAM/MALe_analysis. Raw data are available upon request. RNA-seq counts and associated metadata have been deposited on the Gene Expression Omnibus (GEO) platform for human and myoblast experiments respectively. Accession numbers will be provided upon availability.

## Ethics statement

Ethical approval was granted by Deakin University Human Research Ethics Committee (project number 2021-307).

## Declaration of generative AI and AI-assisted technologies in the writing process

Generative artificial intelligence (AI) tools (ChatGPT) were used to assist with the development and debugging of programming code used in this manuscript. All AI assisted code was critically reviewed and tested by the first author prior to implementation.

## Supplementary material

**Supplementary Figure 1.**
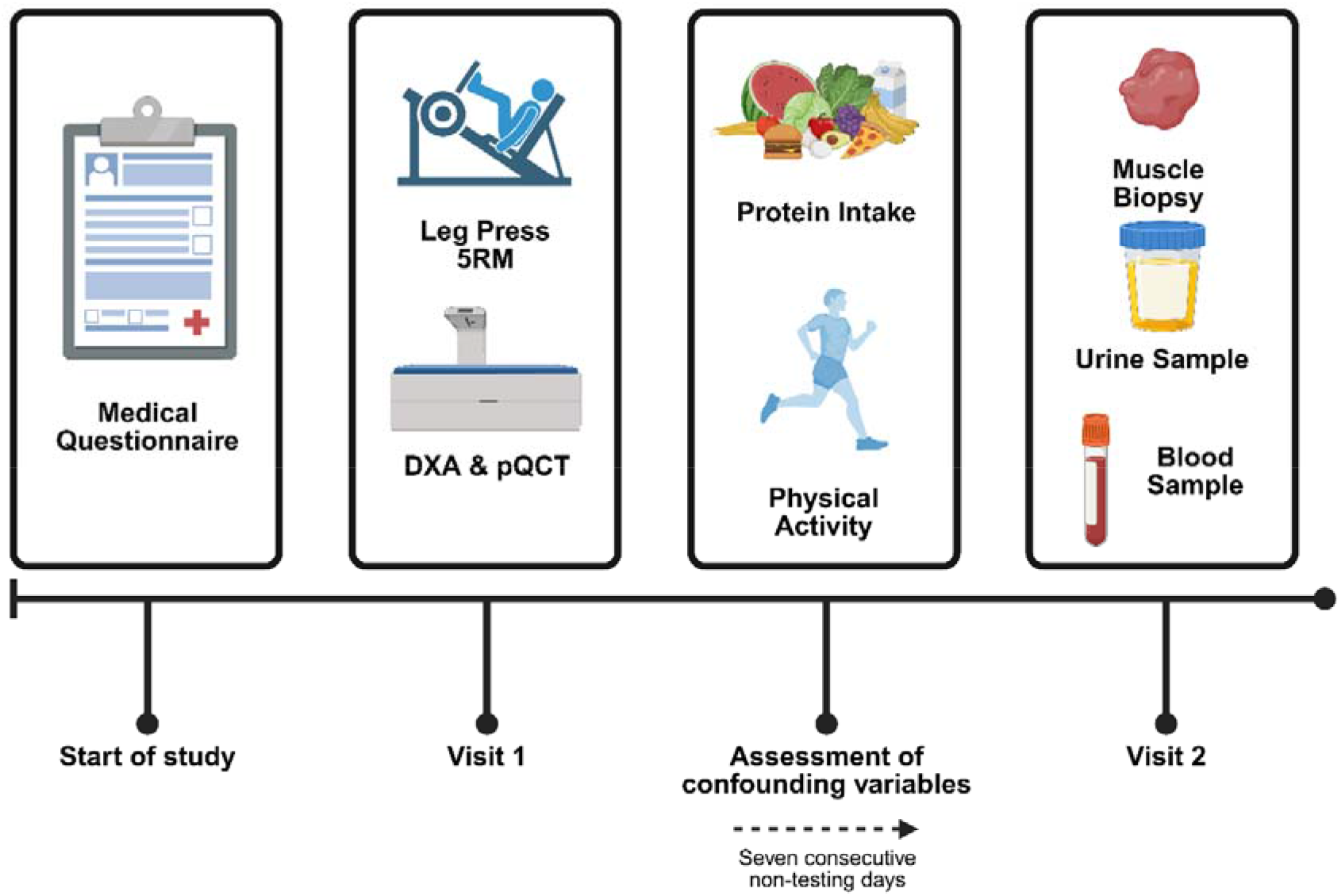
Study design and data collection timeline. 5RM: 5-repetition-maximum; DXA: dual-energy x-ray absorptiometry; pQCT: peripheral quantitative computed tomography.

**Supplementary Figure 2.**
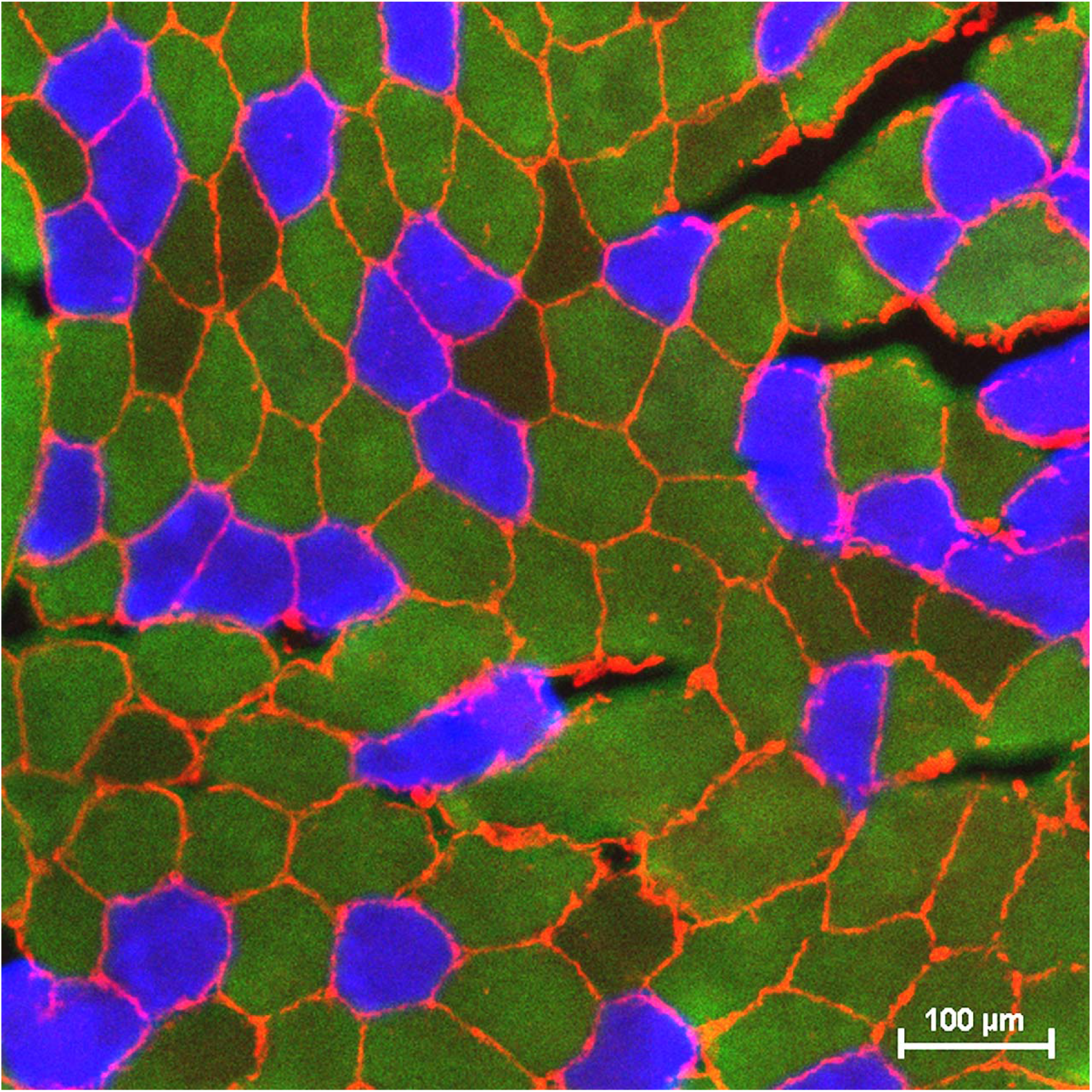
Representative image of muscle fibre typing. Blue represents type I fibres (BA-F8). Green represents type IIa fibres (SC-71). Red represents cell membrane (laminin). Scale bar = 100 µm.

**Supplementary Figure 3.**
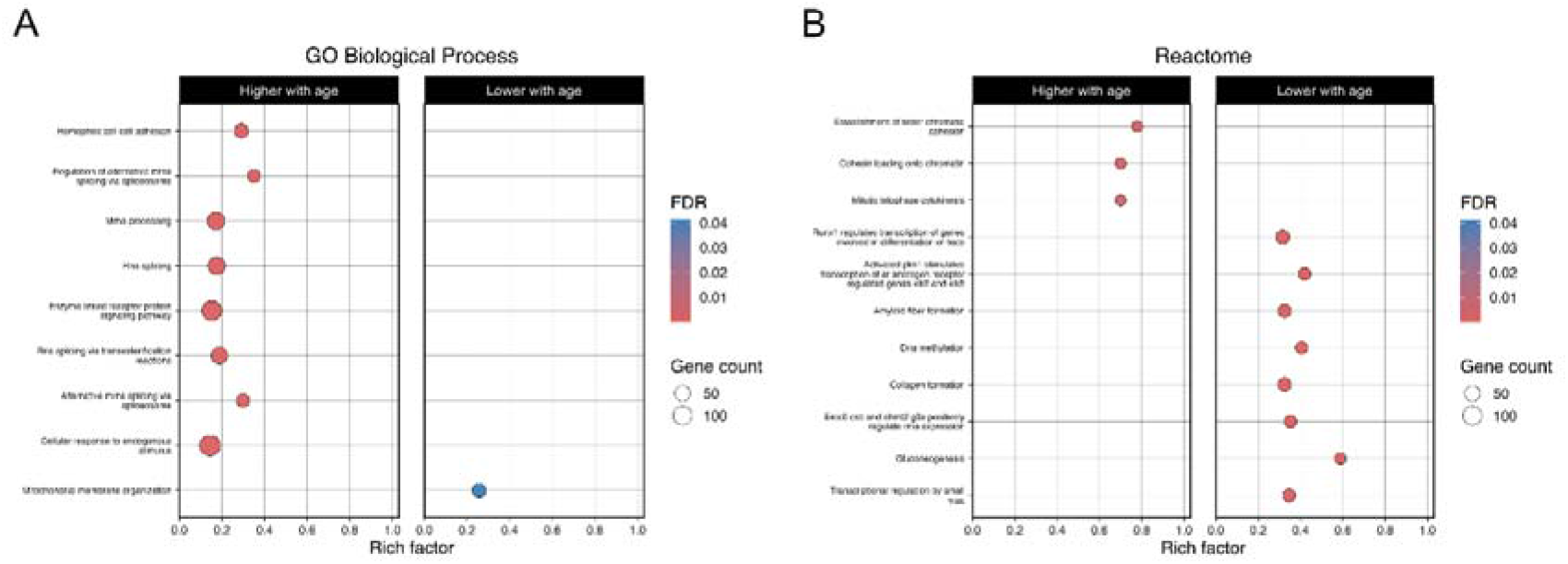
Functional enrichment of age-associated skeletal muscle transcripts. Functional enrichment analysis of genes with significantly different predicted expression between older and younger adult males. Enriched pathways are shown for (A) Gene Ontology (GO) Biological Process and (B) Reactome analyses, stratified according to whether genes showed higher (left) or lower (right) predicted expression in older compared with younger adults. The rich factor represents the proportion of genes within each pathway represented among the age-associated genes. Bubble size indicates the number of genes contributing to each enriched pathway, and colour indicates the false discovery rate (FDR)-adjusted P value, with lower FDR values indicating stronger statistical evidence for enrichment.

**Supplementary Figure 4.**
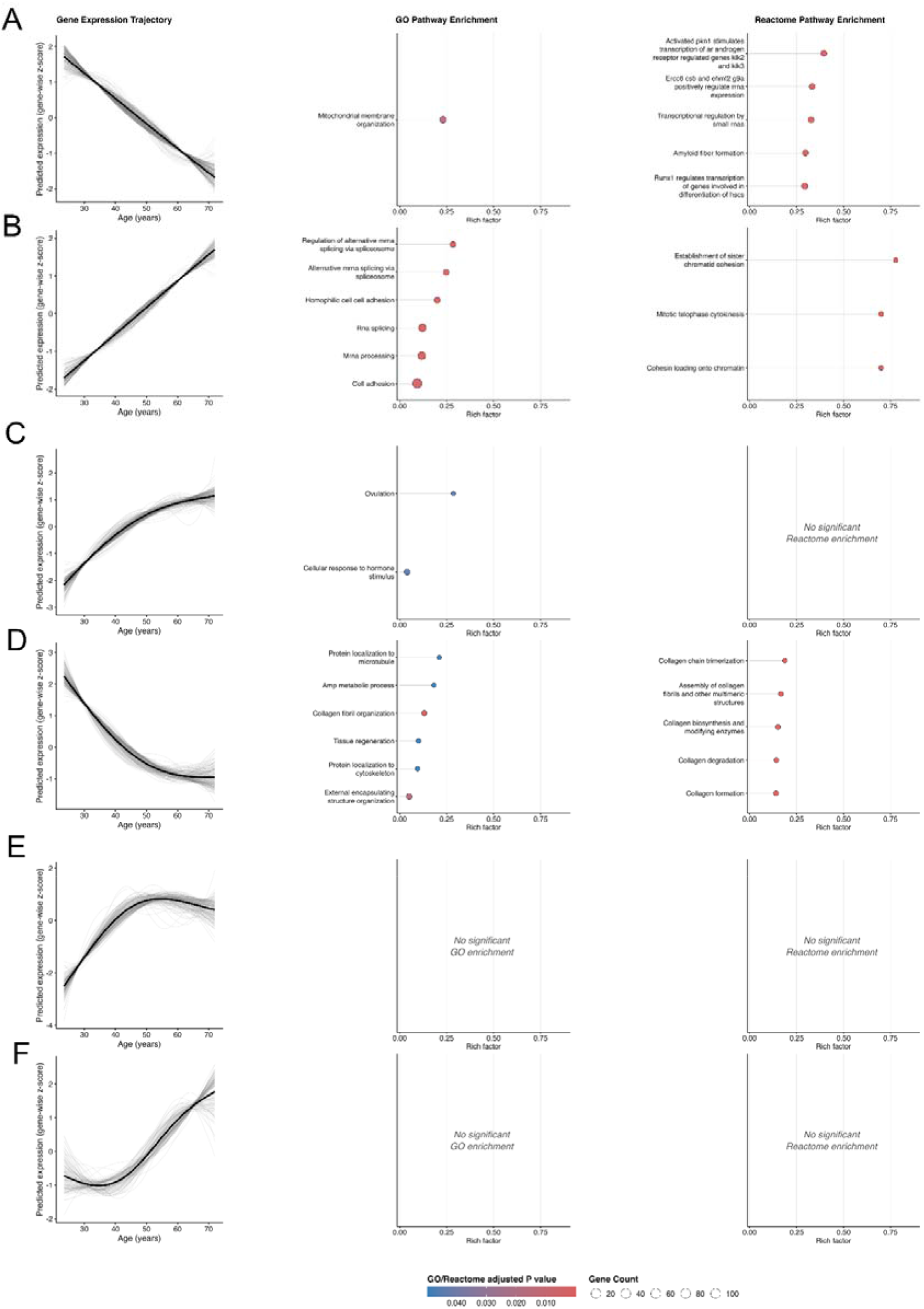
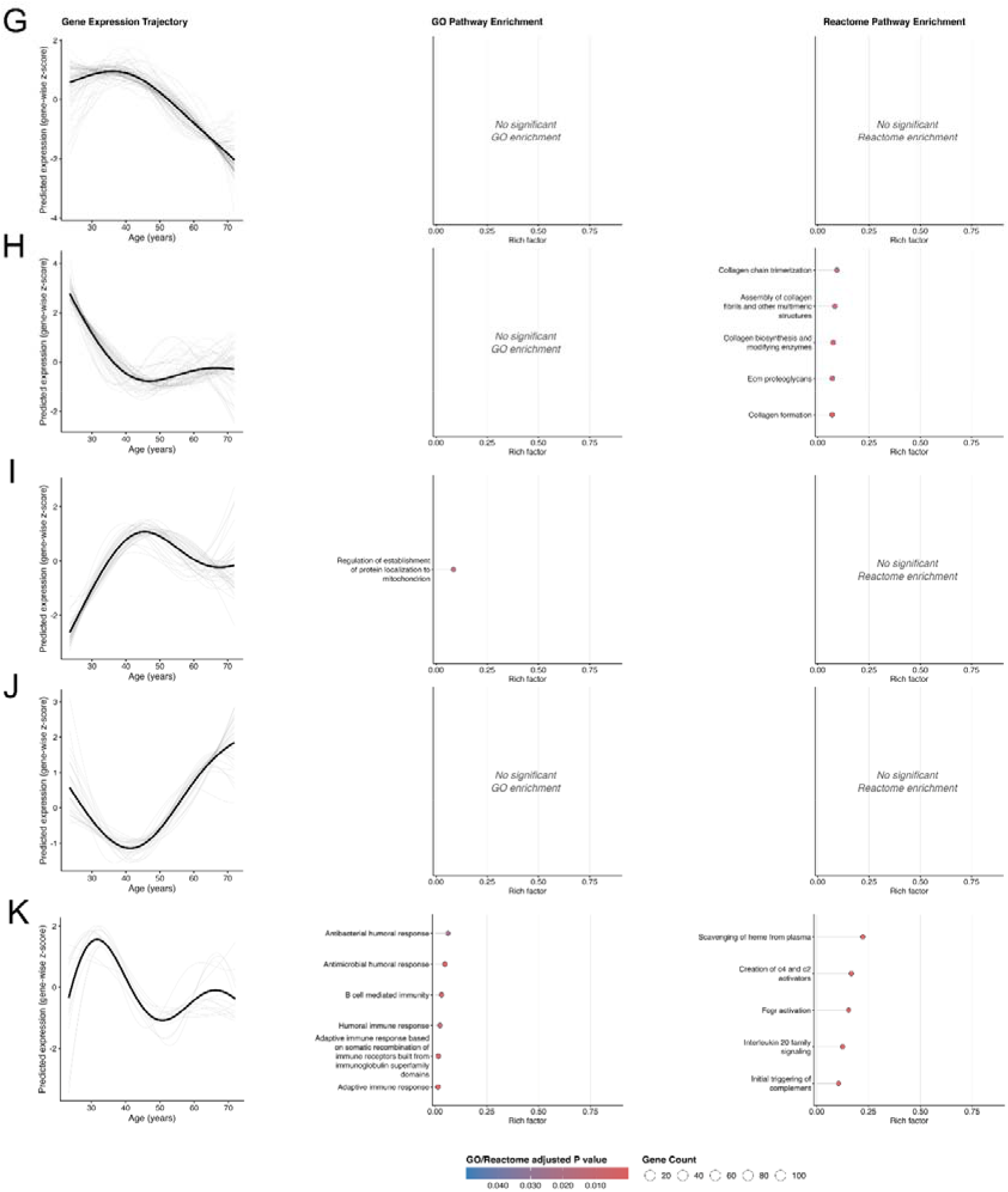
Age-related gene expression trajectories and associated functional pathway enrichment in human skeletal muscle. Genes exhibiting significant age-associated changes in expression were grouped according to their predicted expression trajectories across the adult lifespan. (A–K) represent eleven distinct age-related gene expression trajectories. Left panels show predicted gene expression (gene-wise z-scores) across age for genes assigned to each trajectory; individual gene trajectories are shown in grey and the overall trajectory in black. Middle and right panels show significant enrichment of Gene Ontology (GO) Biological Process and Reactome pathways, respectively, for genes within each trajectory. The rich factor represents the proportion of genes within each pathway represented in the corresponding trajectory. Bubble size indicates the number of genes contributing to each enriched pathway, and colour indicates the FDR-adjusted P value, with lower values indicating stronger evidence of enrichment. Panels without pathways meeting the significance threshold are indicated as having no significant GO or Reactome enrichment.

**Supplementary Figure 5.**
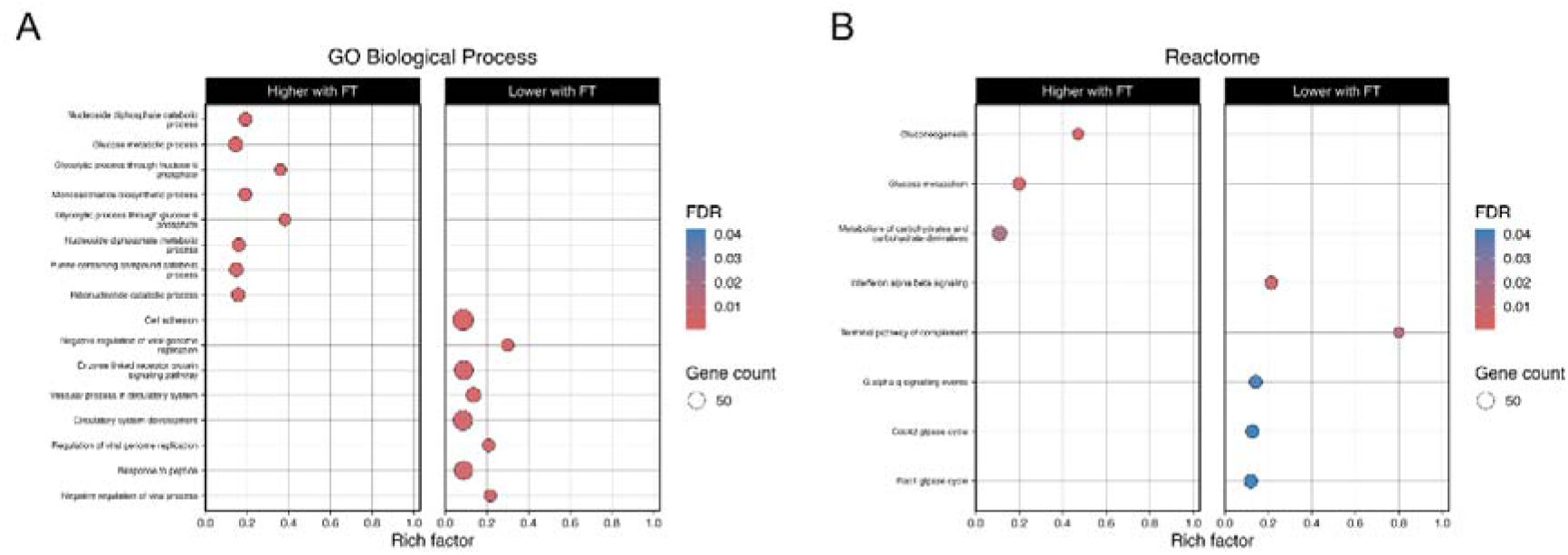
Functional enrichment of free testosterone-associated skeletal muscle transcripts. Functional enrichment analysis of genes with significantly different predicted expression between older and younger adult males. Enriched pathways are shown for (A) Gene Ontology (GO) Biological Process and (B) Reactome analyses, stratified according to whether genes showed higher (left) or lower (right) predicted expression in older compared with younger adults. The rich factor represents the proportion of genes within each pathway represented among the age-associated genes. Bubble size indicates the number of genes contributing to each enriched pathway, and colour indicates the false discovery rate (FDR)-adjusted P value, with lower FDR values indicating stronger statistical evidence for enrichment.

**Supplementary Figure 6.**
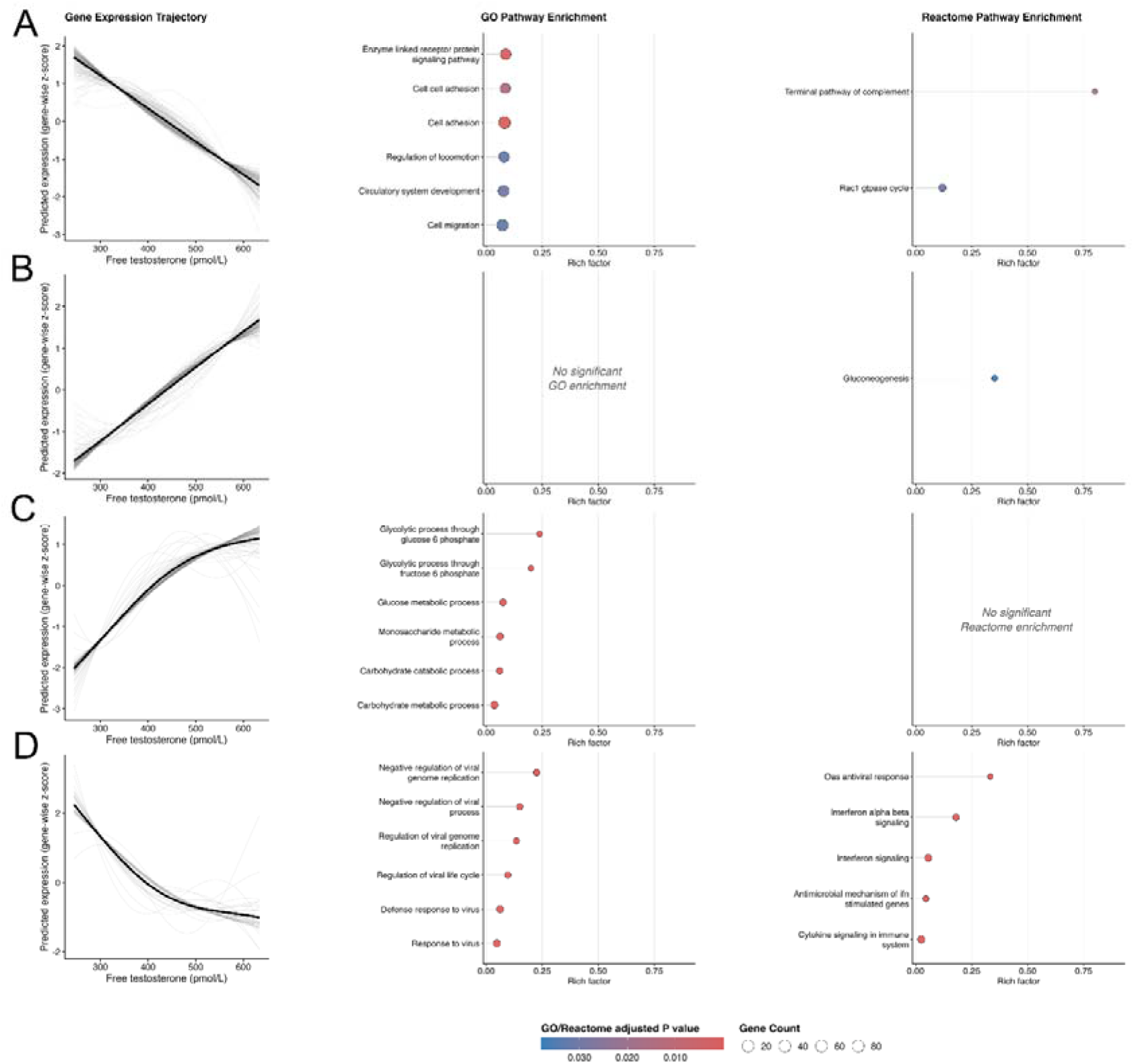
Free testosterone-related gene expression trajectories and associated functional pathway enrichment in human skeletal muscle. Genes exhibiting significant free testosterone-associated changes in expression were grouped according to their predicted expression trajectories across the free testosterone concentration continuum (pmol/L). (A–D) represent four distinct age-related gene expression trajectories. Left panels show predicted gene expression (gene-wise z-scores) across age for genes assigned to each trajectory; individual gene trajectories are shown in grey and the overall trajectory in black. Middle and right panels show significant enrichment of Gene Ontology (GO) Biological Process and Reactome pathways, respectively, for genes within each trajectory. The rich factor represents the proportion of genes within each pathway represented in the corresponding trajectory. Bubble size indicates the number of genes contributing to each enriched pathway, and colour indicates the FDR-adjusted P value, with lower values indicating stronger evidence of enrichment. Panels without pathways meeting the significance threshold are indicated as having no significant GO or Reactome enrichment.

**Supplementary Figure 7.**
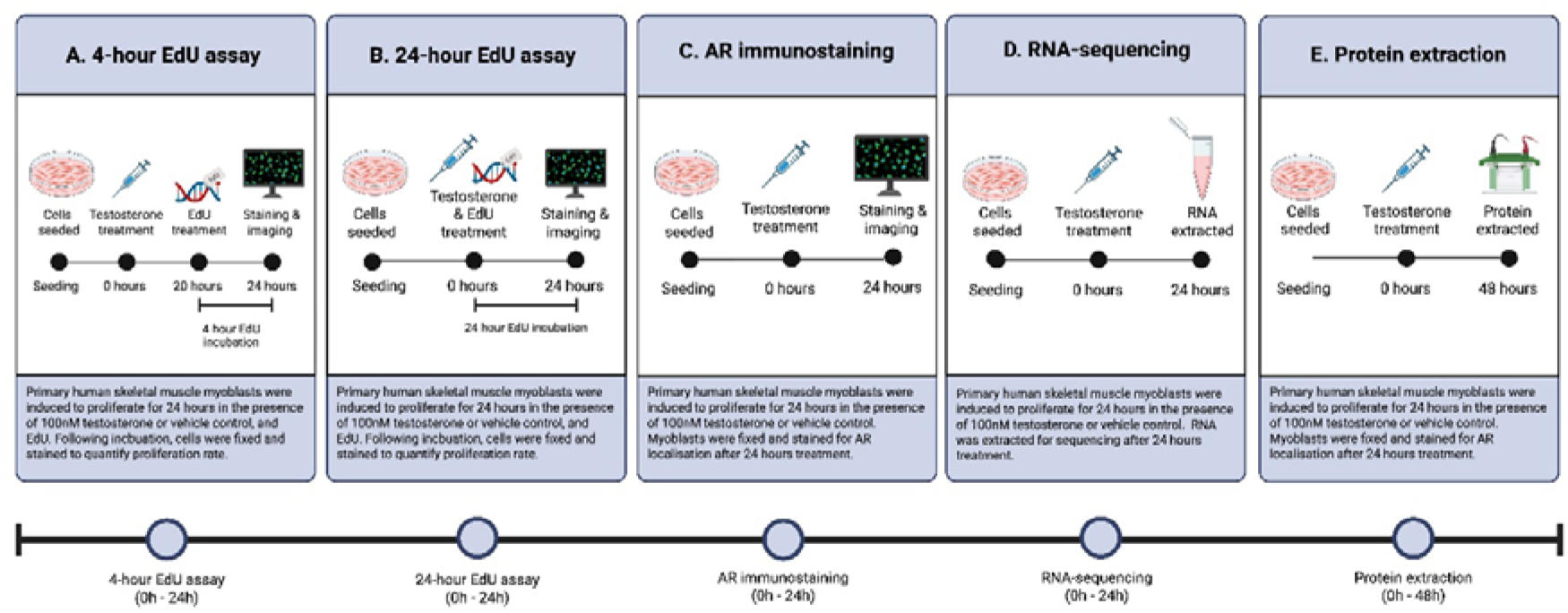
Experimental design for assessing testosterone responses in primary human skeletal muscle myoblasts. Primary human skeletal muscle myoblasts were treated with 100 nM testosterone or vehicle control and assessed using four experimental protocols. (A) Four-hour EdU assay: cells were treated for 24 h, with EdU added for the final 4 h (20–24 h), followed by fixation, staining and imaging to quantify proliferation. (B) Twenty-four-hour EdU assay: EdU was added concurrently with testosterone or vehicle treatment for 24 h, followed by fixation, staining and imaging to quantify proliferation. (C) Androgen receptor (AR) localisation: cells were treated for 24 h before fixation and immunostaining to assess AR localisation. (D) Protein analysis: cells were treated for 48 h before protein extraction for assessment of AR abundance. The lower timeline summarises the timing of each experimental protocol relative to treatment initiation (0 h).

**Supplementary Figure 8.**
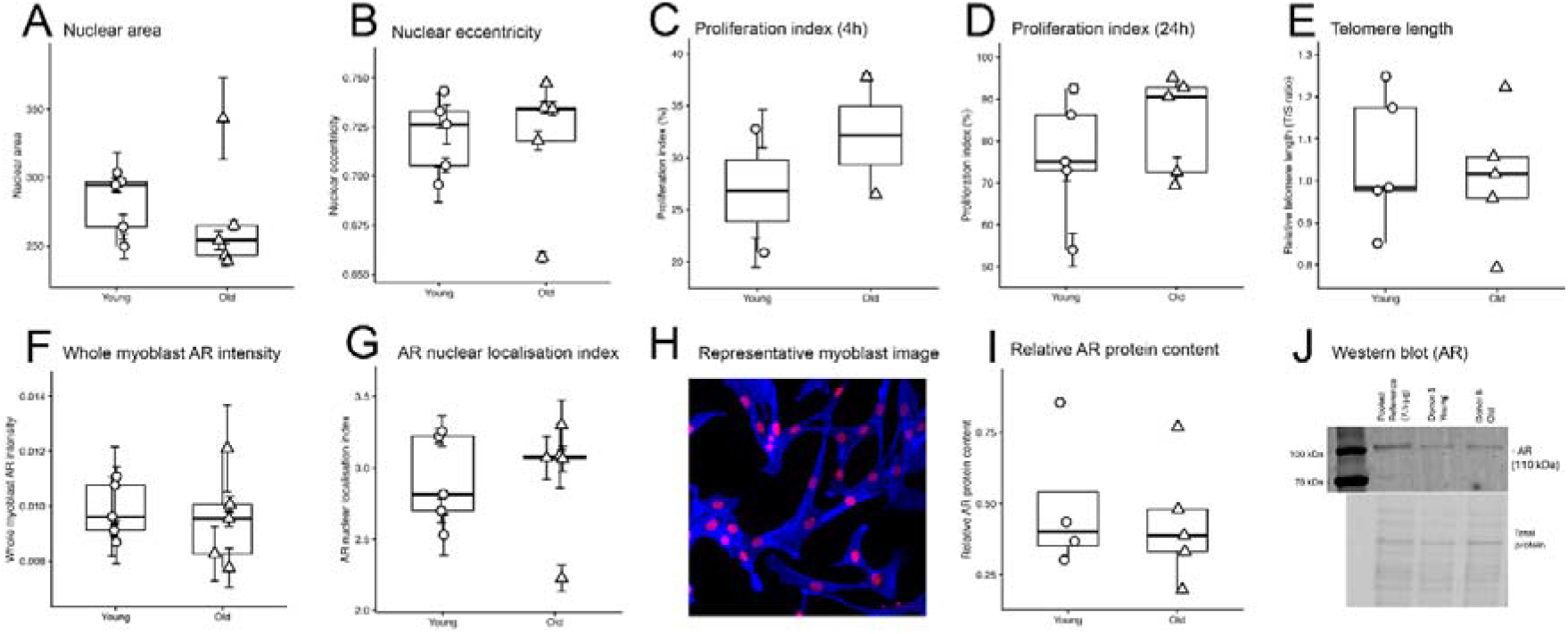
Baseline morphological, proliferative and androgen receptor characteristics of primary human skeletal muscle myoblasts from young and older male donors. Primary human skeletal muscle myoblasts derived from young and older adult males were compared for (A) nuclear area, (B) nuclear eccentricity, (C–D) proliferation index assessed using EdU incorporation, (E) relative telomere length, (F) whole-myoblast androgen receptor (AR) intensity, and (G) AR nuclear localisation index. (H) Representative immunofluorescence image of cultured myoblasts used for assessment of AR expression and localisation. (I) Relative AR protein content assessed by Western blot, with (J) representative AR Western blot and corresponding total-protein image. Box plots show the distribution of individual donor values, with circles representing young donors and triangles representing older donors.

**Supplementary Figure 9.**
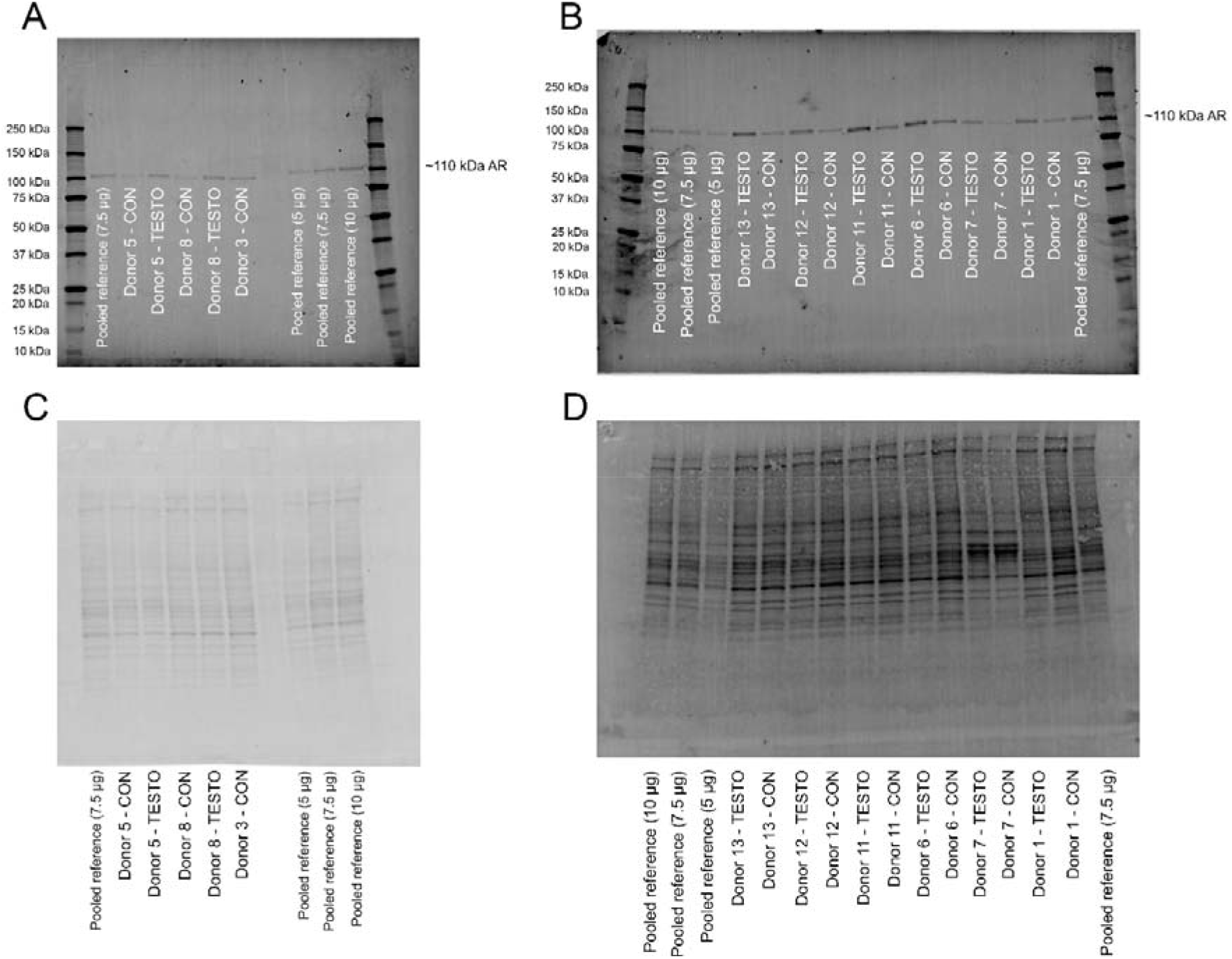
Complete Western Blot data.

**SUPPLEMENTARY FIGURE 10.**
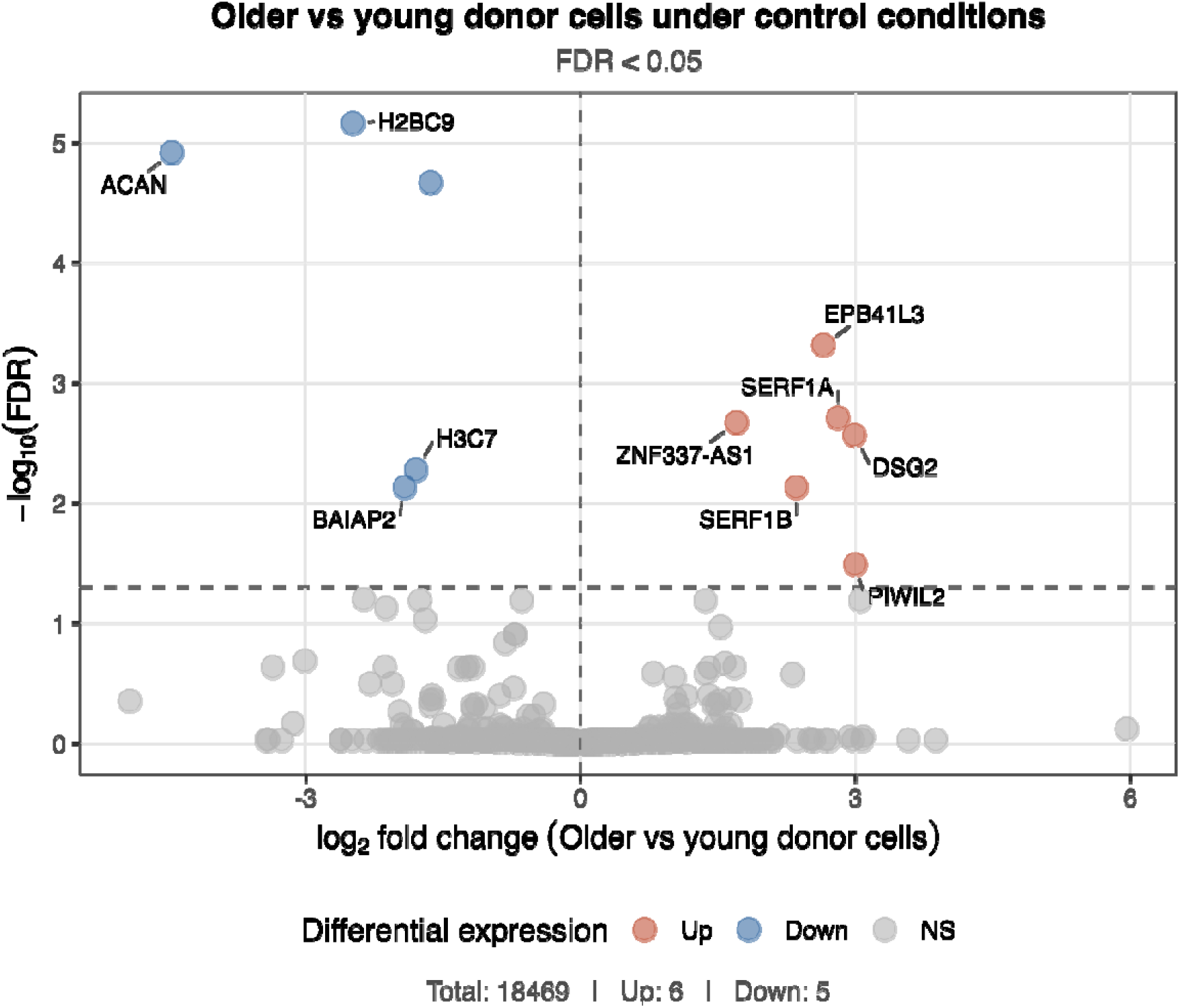
Effect of donor age on the primary human myoblast transcriptome. Volcano plot showing differential gene expression between primary human myoblasts derived from aged (N = 5) and young (N = 5) donors under control conditions. The x-axis represents log fold change (aged/young) and the y-axis represents −log adjusted *P* value. Significantly upregulated and downregulated transcripts (adjusted *P* < 0.05) are shown in red and blue, respectively, with non-significant transcripts shown in grey.

**SUPPLEMENTARY FIGURE 11.**
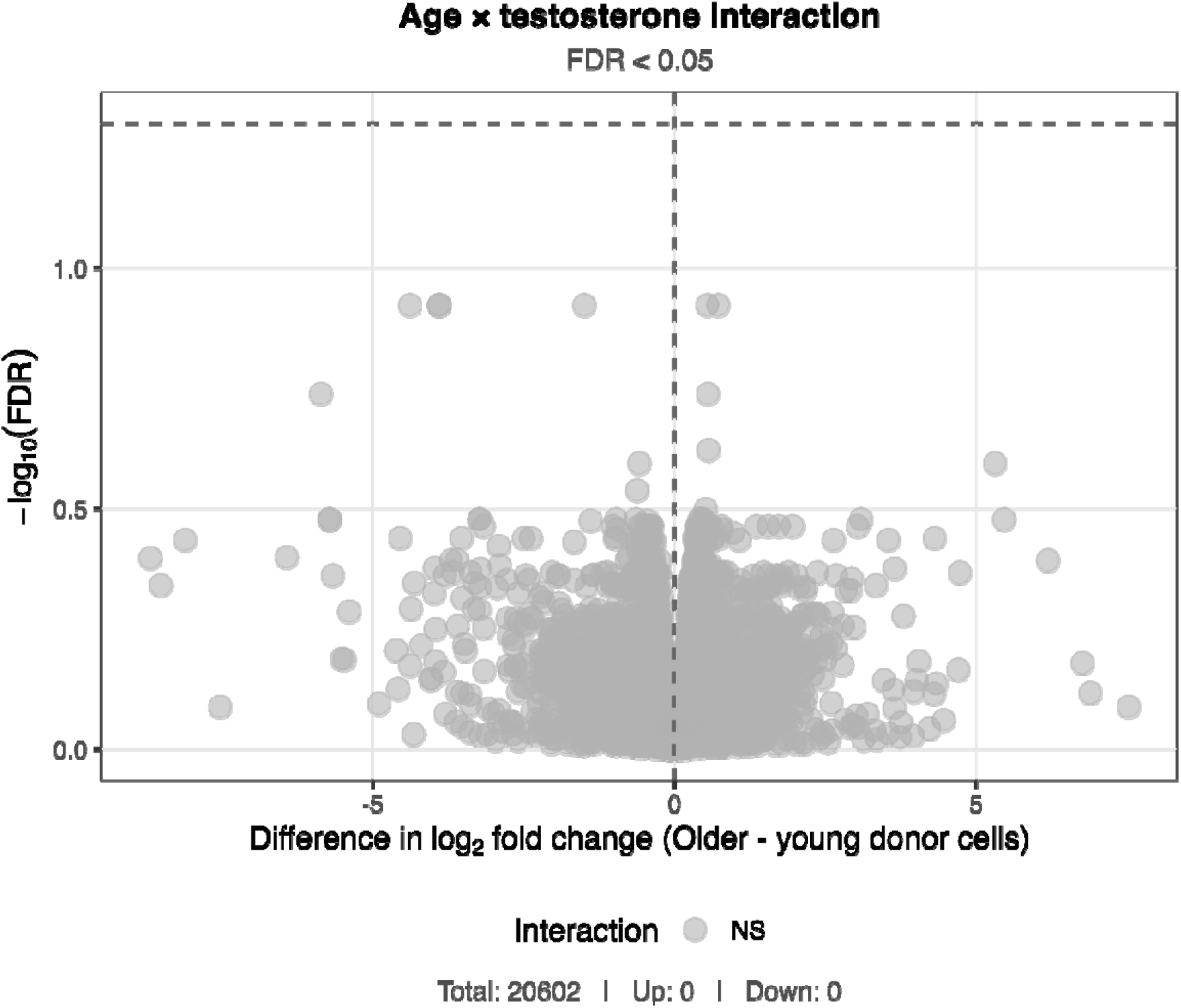
Interaction between donor age and 24hrs 100nM testosterone treatment on the primary human myoblast transcriptome. Volcano plot showing the age × testosterone treatment interaction effect on gene expression in primary human myoblasts derived from aged (N = 5) and young (N = 5) donors. The x-axis represents the log fold change in the transcriptional response to testosterone treatment in myoblasts derived from aged relative to young donors, and the y-axis represents −log adjusted *P* value for the interaction effect.

**SUPPLEMENTARY TABLE 1.** Training status thresholds used to categorize exercise modalities.

| Training status | Sessions per week | Minutes per week |
| --- | --- | --- |
| Resistance trained | ≥ 2 | ≥ 60 |
| Endurance trained | ≥ 3 | ≥ 150 |
| HIIT trained | ≥ 2 | ≥ 40 |
| Body weight | ≥ 2 | - |

**SUPPLEMENTARY TABLE 2.**
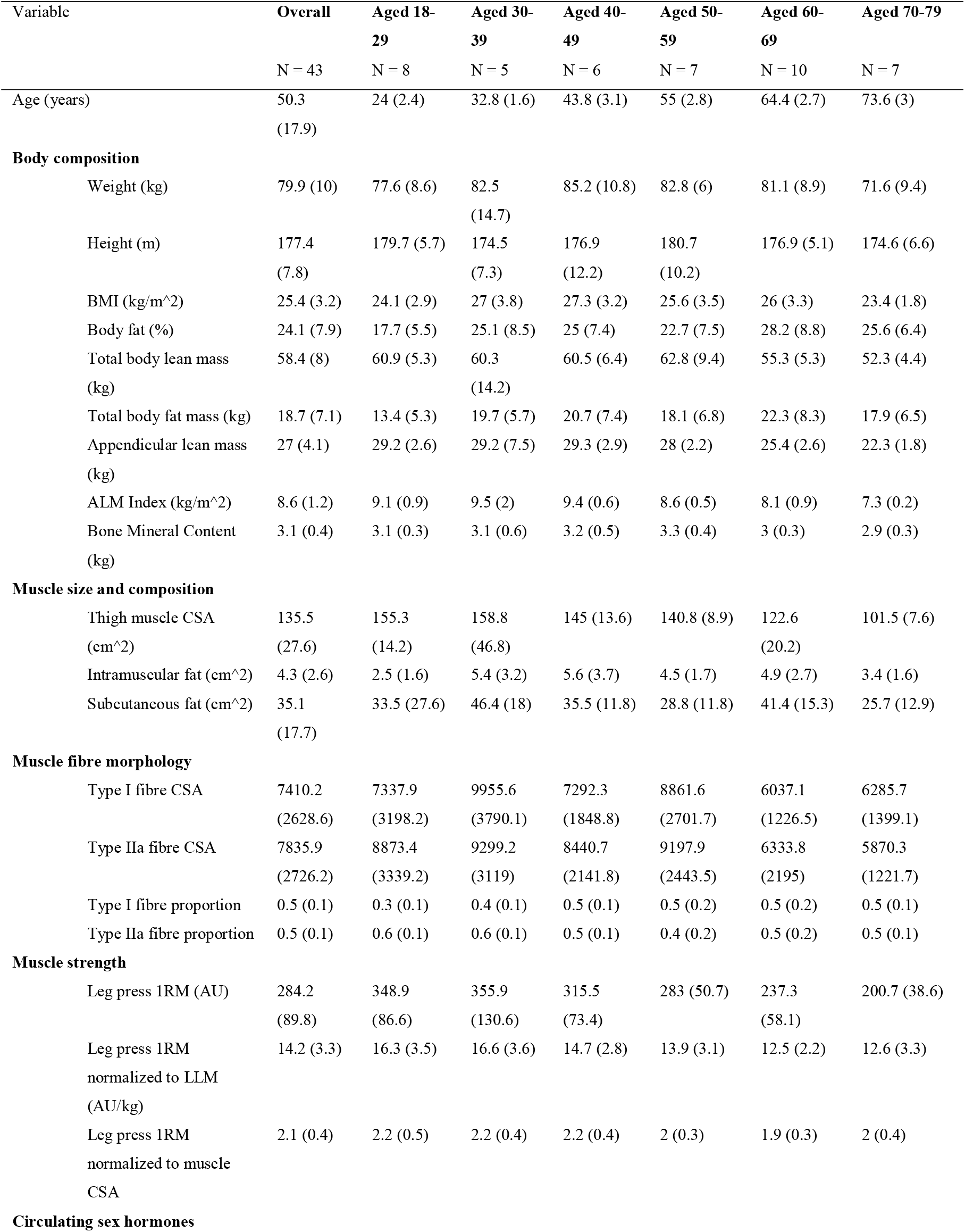
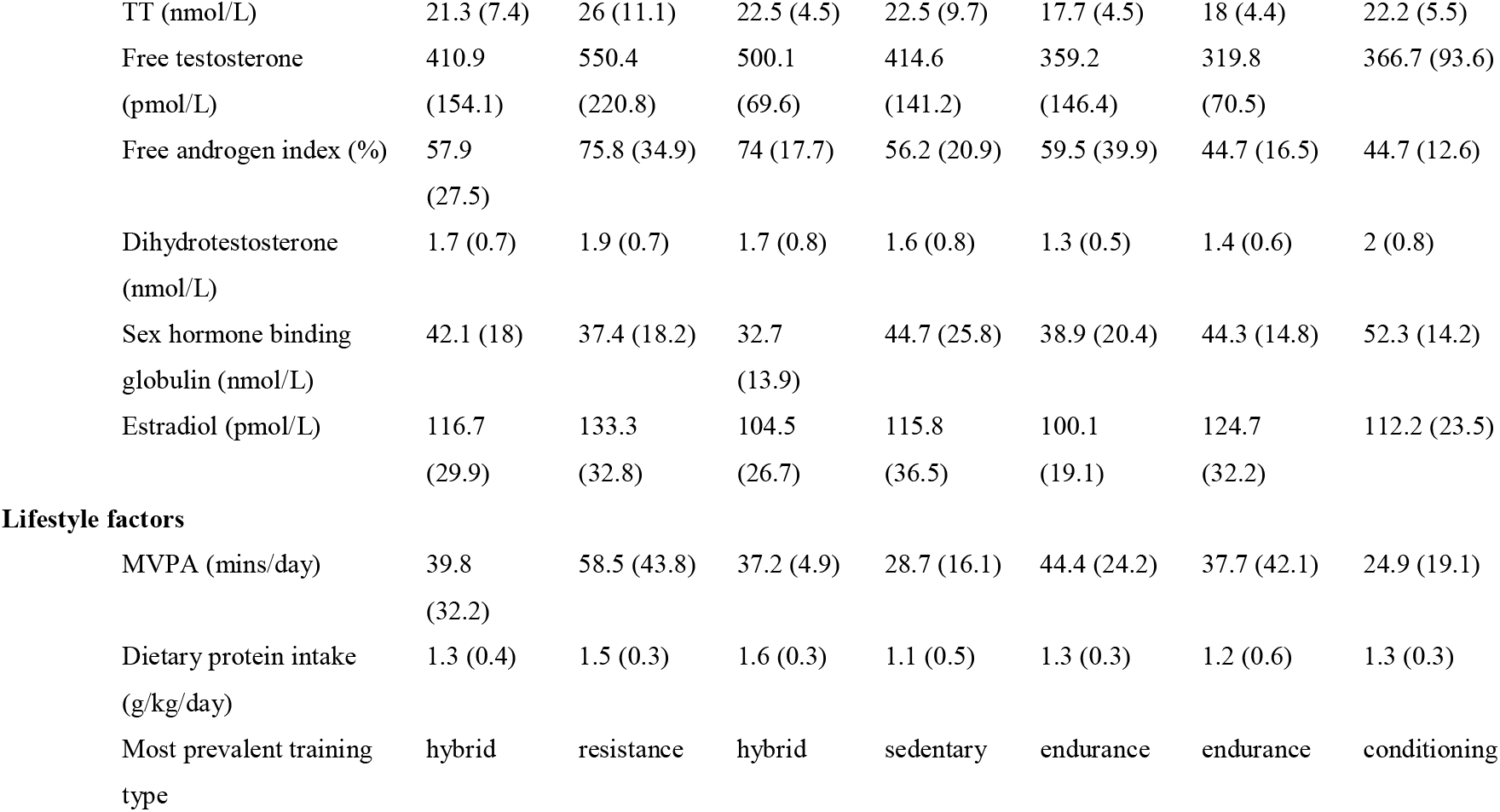
Fibre typing cohort participant characteristics. BMI: Body mass index; ALM: Appendicular lean mass; ALMI: Appendicular lean mass index; CSA: Cross-sectional-area; 1RM: One-repetition-maximum; e1RM: Estimated one-repetition maximum; MVPA: Moderate-vigorous physical activity.

**SUPPLEMENTARY TABLE 3.**
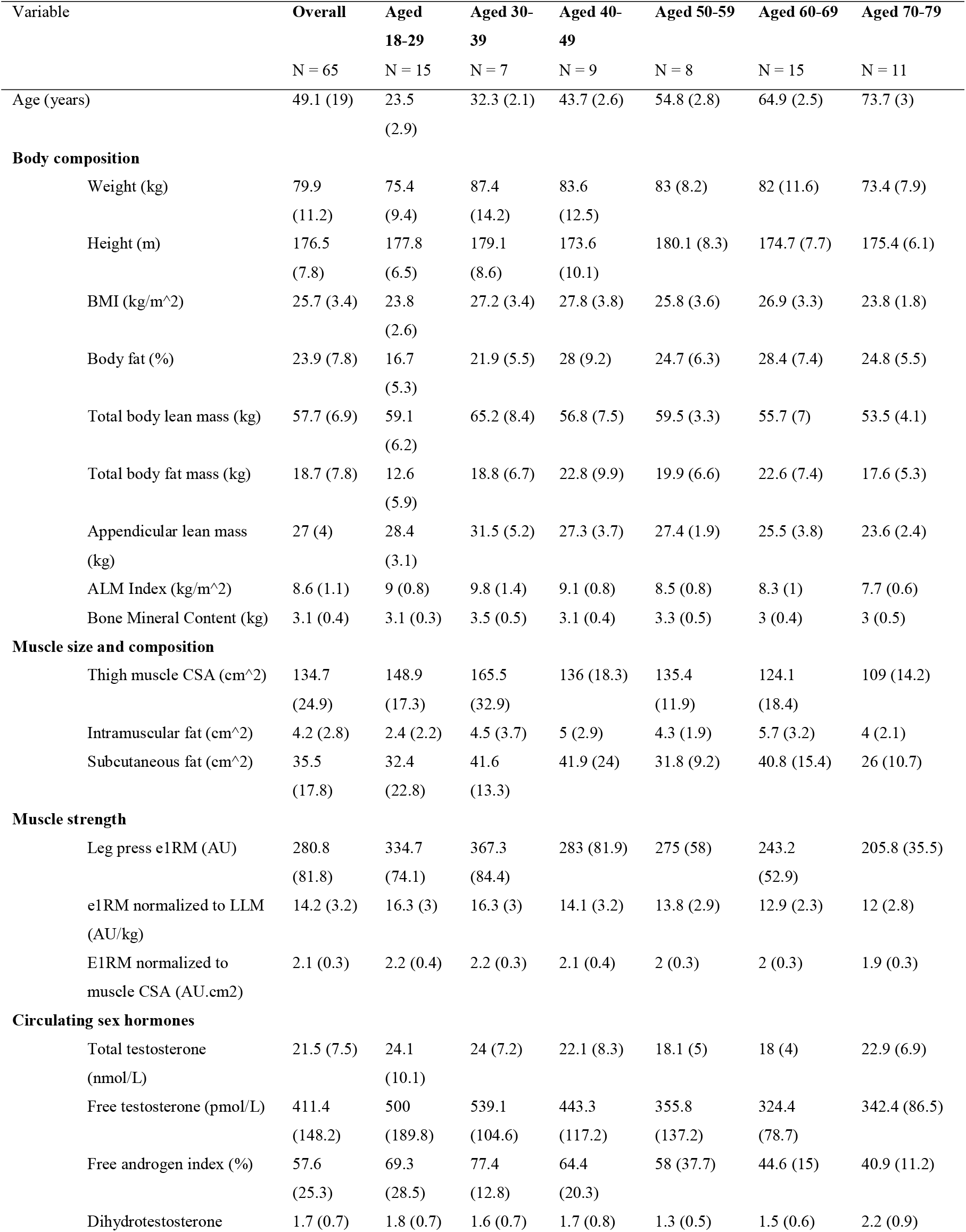
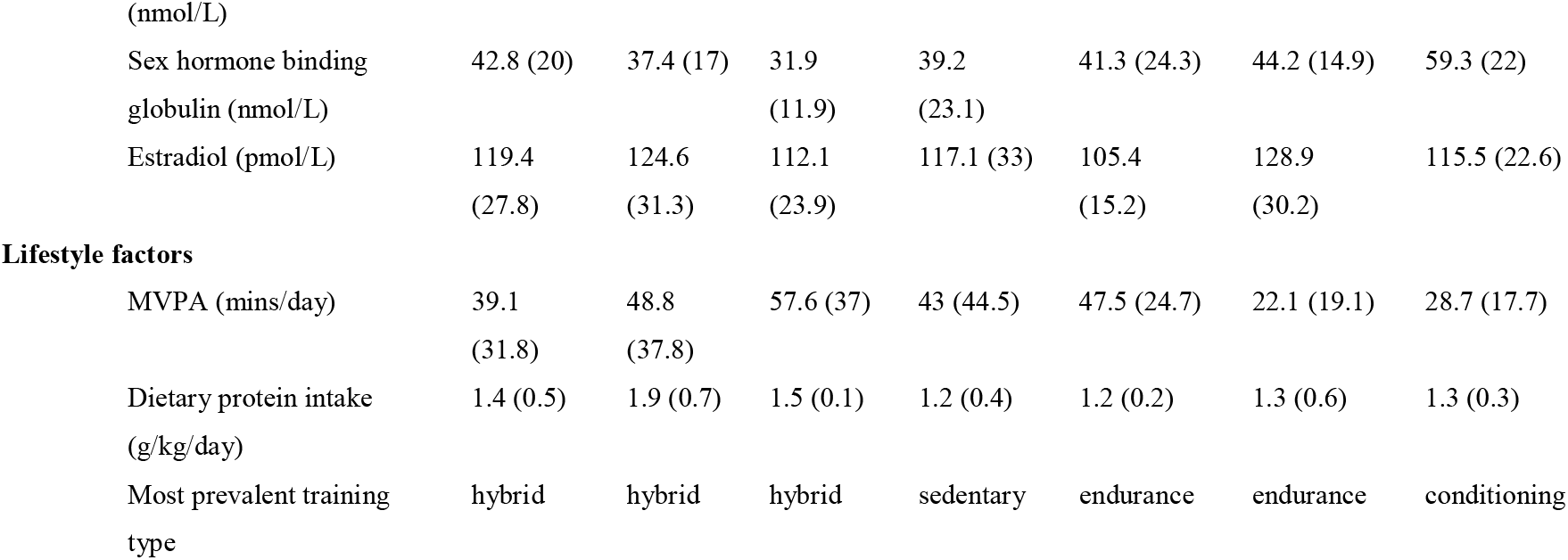
RNA-sequencing cohort participant characteristics. BMI: Body mass index; ALM: Appendicular lean mass; ALMI: Appendicular lean mass index; CSA: Cross-sectional-area; 1RM: One-repetition-maximum; e1RM: Estimated one-repetition maximum; MVPA: Moderate-vigorous physical activity.

**SUPPLEMENTARY TABLE 4.**
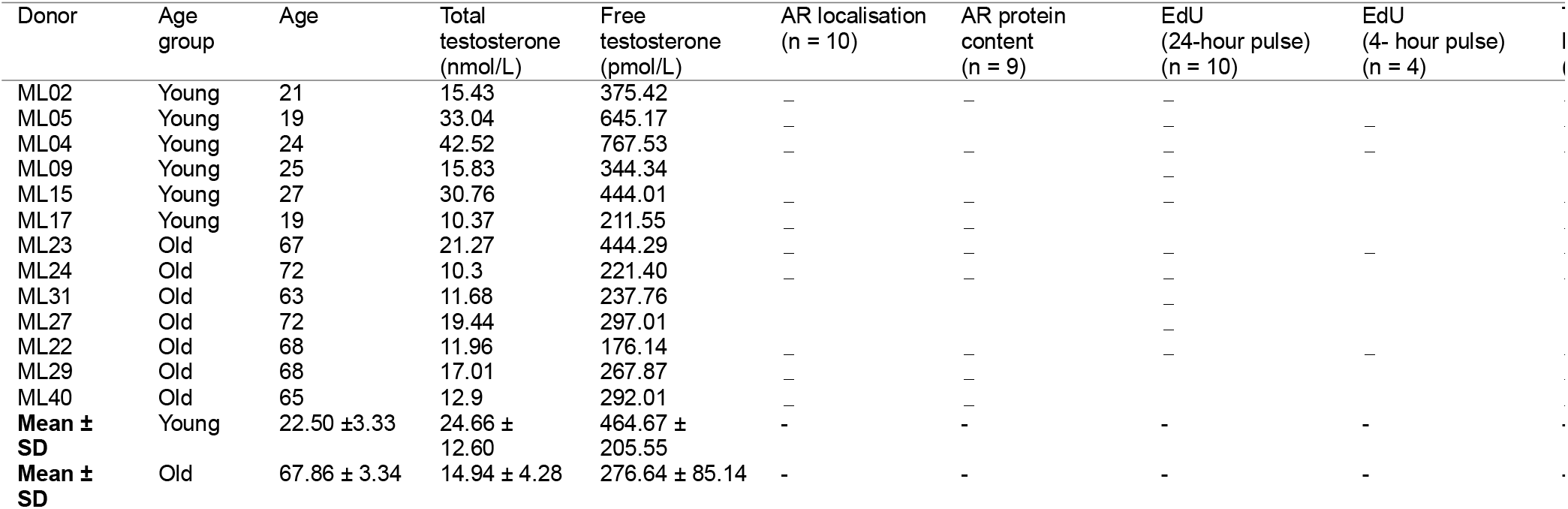
Myoblast donor characteristics.

