## Supplementary Methods for "Molecular responsiveness to testosterone is uncoupled from skeletal muscle mass and strength in males aged 18 - 80 years"

***Western blotting***

Protein concentration of each sample was quantified via the Pierce Bicinchoninic Acid (BCA) assay (cat no. 23225, Thermo Fisher Scientific). Samples were mixed with Laemmli buffer and heated to 95 degrees Celsius for 5 min. 7.5 micrograms of denatured protein was loaded into a 4-20% gradient protein gel (cat no. 5678094, Criterion TGX Stain-Free Precast Gel, Bio-Rad, Gladesville, NSW) and separated via electrophoresis. The stain-free gel was UV-activated for 1 minute and imaged for total protein (BioRad Chemi Doc XR+, Hercules, CA, USA) with ImageLab software (BioRad Image Lab v6). Protein was then transferred to a methanol soaked polyvinylidene difluoride (PVDF) membrane (Millipore Immobilon FL 0.45 µm #IPFL00010). Membranes were blocked in blocking buffer (cat no. 927-70001, Li-Cor Intercept Blocking Buffer, Li-Cor Biosciences, Lincoln, NE, USA) for 1 hour. Membranes were incubated overnight at 4°C in AR primary antibody (cat no. 5153, 1:500, rabbit, Cell Signaling Technology) in blocking buffer (1:2 Licor Intercept Blocking buffer, 1:2 PBS-T, 0.10% SDS) and then incubated in secondary antibody (cat no. 5151, 1:1000, anti-rabbit IgG Dylight 800) in blocking buffer (1:2 Licor Intercept Blocking buffer, 1:2 PBS-T, 0.10% SDS) for 1 hour at room temperature. Proteins were exposed using an Odyssey CLx Infrared Imaging System (Li-Cor Biosciences, Lincoln, NE, USA) and protein band densities determined using the Odyssey Image Studio software (Image Studio V5.2, Li-Cor Biosciences, Lincoln, NE, USA). A pooled reference sample was formed from 8.5ug of protein from each sample and loaded onto each gel. AR protein quantification was normalised against sample total protein content and against the pooled sample within each gel. Furthermore, 5, 7.5 and 10 µg protein of the reference sample was loaded to confirm the AR band intensity was within the linear dynamic range (1, 2).

#### **Immunohistochemical staining and imaging (supplementary)**

*Vastus lateralis* muscle samples were immunohistochemically stained to quantify muscle fibre type and size as described in (3). Biopsied tissue was sectioned into 20 µm sections using a cryostat set to -18°C and were mounted on coverslips and stored at -80°C until further use. Prior to staining, sections were thawed for 30 minutes at room temperature and washed with phosphate-buffered saline (PBS). Sections were incubated for one hour in blocking buffer (10% goat serum, cat. no. 16210072, Thermo Fisher Scientific; 1x PBS), then for one hour with a primary antibody cocktail in blocking buffer: BA-F8 (DSHB, University of Iowa, 1:20), SC-71 (DSHB, University of Iowa, 1:50), laminin (Sigma Aldrich, 1:100). Sections were then washed with PBS and incubated concealed from light for one hour with a secondary antibody cocktail in PBS: Goat α-mouse IgG2b Alexa Fluor 647 (cat. no. A-21242, Thermo Fisher Scientific, 1:500), goat α-mouse IgG1 Alexa Fluor 488 (cat. no. A-11001, Thermo Fisher Scientific, 1:500), goat α-rabbit IgG Alexa Fluor 405 (cat. no. A-31556, Thermo Fisher Scientific, 1:500). Coverslips were sealed using Vectashield Vibrance (cat. no. H1700-2, Vector Laboratories, Newark, CA, USA).

Sections were imaged using the Eclipse Ti2 microscope (Nikon, Tokyo, Japan) and NIS-Elements Advanced Research software (Nikon, Tokyo, Japan). Images were taken at 4x objective, 1024x resolution using a 4x line average and 6.2 µs pixel dwell time. Where sections were larger than the imaging field, multiple images were taken and stitched together using the Large Image function in NIS-Elements. Images were analysed using the SMASH (semi-automated muscle analysis using segmentation of histology) toolbox (4) for MATLAB (R2023b, The MathWorks In., Natick, Massachusetts). The cross-sectional area of each individual fibre was quantified and the number and proportion of each fibre type (types I and IIa) calculated. Sections with < 100 fibres were excluded from analysis (5). A representative image is provided in **Supplementary Figure 2.**

***Proliferation index, nuclei size and eccentricity***

Proliferation index was measured using a Click-iT EdU Cell Proliferation kit (cat. No. C10340, Thermo Fisher Scientific). At the onset of treatment, cells were incubated in 1x EdU for 24 hours. Cells were then fixed in 4% PFA/PBS for 15 minutes and permeabilised in 0.5% Triton X-100 (cat. No. A16046-AE, Sigma-Aldrich) / PBS for 20 minutes. Cells were incubated with Click-iT reaction cocktail for 30 minutes at room temperature, then incubated in 1x Hoechst 33342 for 30 minutes at room temperature. A sub-set of cells (n = 2 young adult donors, n = 2 older adult donors) were also incubated in 1x EdU for the final 4 hours of a 24-hour testosterone treatment.

Cells were imaged using the Eclipse Ti2 microscope and NIS-Elements Advanced Research software. Images were obtained at 20x objective and 1024x resolution using a 4x line average and 2.2 µs pixel dwell time. Per cell line, 2x wells were imaged per treatment, with 12x fields imaged per well. Coordinates for each field were systematically randomised, with each well-being divided into a 4x3 grid and coordinates within each grid square randomly generated. Nuclei were segmented using a model trained in CellPose (V3.0) (6). The model was validated through manual identification of nuclei in 10% of each imageset in duplicate. Segmented images were reviewed and manually amended where required. Quantification of proliferation index, nuclei size and nuclei eccentricity was performed using the open-source image analysis software CellProfiler (V4.2.5) and an analysis pipeline developed as part of this study. Segmented nuclei masks were imported into CellProfiler. Nuclei area and eccentricity were measured per nuclei and the proliferation index per field calculated using the following equation:

$$Proliferation index= \frac{Total Nuclei}{{EdU}^{+}Nuclei} \times100$$

#### **AR immunohistochemical staining and imaging**

Fixed cell samples were permeabilised in 0.1% Triton X-100/PBS for 5 minutes and blocked with blocking buffer (3% BSA (cat. No. A9418, Sigma-Aldrich) / PBS) for 40 minutes at room temperature. Cells were then incubated with the primary antibody against the AR (cat. no. 5153, 1:50, Cell Signaling Technology) in blocking buffer for 1 hour at room temperature. Cells were then incubated with AlexaFluor 488 goat anti-rabbit IgG (cat. no. A‐11008, Thermo Fisher Scientific, 1:5000) and AlexaFluor 647 phalloidin (cat. No. A12379, Thermo Fisher Scientific, 1:200) in 1% BSA/PBS for 1 hour at room temperature. Cells were then incubated with DAPI (cat. No. 62248, Thermo Fisher Scientific, 1:1000) in PBS for 10 minutes. Images were obtained at 40x objective and 1024x resolution using a 4x line average and 6.2 µs pixel dwell time. Per cell line, 2x wells were imaged per treatment, with 12x fields imaged per well as described in section 2.2.5. Nuclei were segmented using a model trained in CellPose (V3.0) (6). Myoblasts were manually segmented using Cellpose. Myoblasts were defined as mononucleated cells expressing actin. Segmented myoblast and nuclei masks were imported into CellProfiler. Mean cell AR intensity was measured using the myoblast masks. This cell-level measure represents the concentration of AR-associated fluorescence within individual myoblasts, and was distinguished from western blotting, which measured relative AR protein abundance in pooled whole-cell lysates. To quantify AR nuclear localisation, a non-nuclear mask was generated for each myoblast excluding the nuclear-stained region. Mean AR intensity was measured within each nucleus and within each non-nuclear region, and the AR nuclear localisation index calculated using the following equation:

$$AR nuclear localisation index=\frac{Mean nuclear AR intensity}{Mean non nuclear AR intensity}$$

#### **Telomere length**

DNA was extracted from cell lysates using an AllPrep DNA/RNA/miRNA Universal Kit (cat no. 80224, Qiagen, Hilden, Germany). DNA quality and quantity were assessed using a TapeStation system (Agilent Technologies). Telomere length was measured via quantitative PCR (qPCR) as described in (7). Primer sequences were as follows: Telomere Forward Primer Sequence CGGTTTGTTTGGGTTTGGGTTTGGGTTTGGGTTTGGGTT; Reverse Primer Sequence GCTTGCCTTACCCTTACCCTTACCCTTACCCTTACCCT. 36B4 Forward Primer Sequence CAGCAAGTGGGAAGGTGTAATCC; Reverse Primer Sequence CCCATTCTATCATCAACGGGTACAA. Relative telomere length was expressed as the telomere (T) to single copy gene (S) ratio (T/S ratio), relative to the median sample T/S ratio.

***Transcriptomic analysis (human tissue)***

Transcriptomic analysis was performed in RStudio (Version 2025.09.2+418). Raw gene count matrices and sample data were imported (n = 65) and genes with at least 10 counts in at least 20% of samples retained. A total of 17522 genes were retained for analyses. Gene identifiers were annotated using the org.Hs.eg.db annotation database and *AnnotationDbi* package (8). The *DESeq2* (9) package was used to estimate library-size factors and gene-specific negative-binomial dispersion parameters. Expression of each gene was modelled independently using negative-binomial generalised additive models (GAMs) with restricted maximum likelihood (REML) smoothing parameters. Separate models examined the association between gene expression and age or hormone concentrations. Models included sequencing batch as a covariate, a smooth term for either age or hormone concentration and a library size offset derived from *DESeq2* size factors. Age and hormones were modelled as penalised regression splines with a basis dimension of k = 6 using the *mgcv* package. Adequacy of spline basis dimension was assessed using the *k*-index, with basis dimensions considered adequate if *k*-index > 0.9 and the associated *k-*check *P* > 0.05. Genes failing this assessment were refit with a basis dimension of k = 8. Diagnostic statistics were retained for quality assessment but were not used as exclusion criteria. Estimated degrees of freedom (EDF), test statistics and *P* values were extracted from each GAM. *P* values were adjusted across genes using the Benjamini-Hochberg procedure. Models which failed to fit or had non-finite *P* values were excluded prior to FDR correction. Gene associations with an FDR < 0.05 were considered statistically significant. Gene associations were classified according to EDF. Gene associations with an EDF < 1.2 were classified as approximately linear relationships.

Log_2_ endpoint effect sizes were calculated as the difference between predicted expression at the 90^th^ and 10^th^ percentiles of the observed age or hormone distribution on the model link scale, divided by log(2). Additionally, the maximum-to-minimum predicted expression difference across the age or hormone continuum was calculated to characterise the magnitude of nonlinear trajectories. Significantly associated genes were visualised as heatmaps using the *ggplot2* package (10).

#### **Clustering analysis**

For each separate age and hormone model, FDR significantly associated genes were clustered based on expression trajectory across the respective age or hormone continuum. Clusters were identified based on temporal expression pattern, rather than absolute magnitude of expression. Predicted expression trajectories were standardised using z-score transformation. Trajectory clusters were identified using pairwise Euclidean distances and hierarchical clustering using Ward.D2 linkage in the *dynamicTreeCut* package (11), with a minimum cluster size of 10 genes and moderate tree-splitting sensitivity (deepSplit = 4). Results were visualised as gene expression trajectory cluster plots using the *ggplot2* R package (10).

***Functional analysis***

Global directional and trajectory cluster specific functional enrichment analysis was performed using the Gene Ontology Biological Process and Reactome databases via the *msigdbr* package (12). Over-representation analysis was performed using hypergeometric testing of clusters containing at least 10 genes using the *clusterProfiler* package (13). The analysis-specific background comprised genes which passed expression filtering, had valid gene-specific dispersion estimates and produced successfully fitted GAMs with finite *P* values. Enrichment *P* values were adjusted using the Benjamini-Hochberg correction, with significance set at FDR < 0.05. Results were visualised as directional over-representation analysis (ORA) and cluster specific dot plots using *ggplot2* and *clusterProfiler* R packages (10, 13).

***Age x hormone interaction analyses***

Age x hormone interaction analyses were performed separately for each hormone. Genes were modelled independently using GAMs as described above. Models included sequencing batch as a covariate, a smooth term for age, a smooth term for hormone concentration, a tensor-product interaction smooth for age x hormone and a library size offset derived from *DESeq2* size factors. *P* values for the interaction were adjusted across genes using the Benjamini-Hochberg correction. The interaction effect was summarised as the predicted hormone effect across the 10^th^-90^th^ percentiles of the age continuum, where the hormone effect at each age was defined as the delta between predicted gene expression at high (90^th^ percentile) hormone concentration and predicted gene expression at low (10^th^ percentile) hormone concentration.

***Magnetic Activated Cell Sorting (MACS)***

Once the cells had reached 70-80% confluency, myogenic satellite cells were purified using Magnetic Activated Cell Sorting (MACS) columns with anti-CD56^+^ microbeads (cat. no. 130‐050‐401, Miltenyi Biotec, Bergisch Gladbach, Germany). Cells were trypsinised, pelleted and resuspended in MACS separation buffer (cat. No. 130-091-221, Miltenyi Biotec, Bergisch Gladbach, Germany). Cells were then incubated for 15 minutes at 2-8°C with anti-CD56^+^ microbeads, before being pelleted and resuspended in fresh MACS separation buffer. The cell suspension was then applied to MACS separator columns in a MACS separator and the flow through discarded. Myoblasts were flushed from the MACs columns with separation buffer into a flask pre-coated with ECM and filled with pre-warmed proliferation media. Cells were incubated at 37°C and 5% CO_2_ with proliferation media changed every 48 hours.

***Transcriptomic analysis (cells)***

RNA was isolated and sequenced as described above. Transcriptomic analysis was performed using the *DESeq2* package (9). Raw gene count matrices and sample data were imported and genes with at least 10 counts in at least five samples were retained. Gene symbols and biotypes were obtained from the Ensembl release 115 annotation. Ensembl gene identifiers were mapped to Entrez gene identifiers using the org.Hs.eg.db annotation database and *AnnotationDbi* (8) package. The *DESeq2* package was used to estimate library-size factors and gene-specific negative-binomial dispersion parameters.

Differential expression was modelled using negative-binomial generalised linear models. Testosterone effects were modelled using donor age group, donor nested within age group and age-group-specific treatment effects. Donor terms encompassed the paired control and testosterone treated samples from each donor. Contrasts were used to estimate testosterone treatment effects within each donor age group and the overall treatment effect, calculated as the average of the two age-group effects. An additional contrast tested the age x treatment interaction. The age x treatment interaction was defined as the testosterone-versus-control log_2_ fold change in older donors minus the log_2_ fold change in young donors. Age group differences were analysed separately within control and testosterone-treated samples.

Differential expression was assessed using two-sided Wald tests against a log_2_ fold change of zero. *P* values were adjusted using the Benjamini-Hochberg correction. Genes with an FDR < 0.05 were considered significantly differentially expressed. Log_2_ fold change estimates were reported without shrinkage, where a positive change indicated higher expression with testosterone for treatment effects, and higher expression in older donors for age group effects. For the interaction effect, positive values indicated a more positive testosterone response in older compared to young donor cells.

Gene set enrichment analysis was performed separately for each contrast using the Reactome database and *ReactomePA* package (14). Genes were ranked by their signed *DESeq2* Wald statistic, including those which were not differentially expressed. Where multiple Ensembl identifiers mapped to the same Entrez identifier, the gene with the largest absolute Wald statistic was retained. Analysis was restricted to gene sets containing 10-500 genes represented in the ranked list. Enrichment *P* values and normalised enrichment scores (NES) were estimated using the *fgsea* package (15), with an eps of 0. Enrichment *P* values were adjusted using the Benjamini-Hochberg correction, with significance set at FDR < 0.05. Differential expression and enrichment results were visualised using volcano plots and directional enrichment plots.

***Statistical analysis (cells)***

Immunohistochemical measures in proliferating myoblasts were analysed using linear mixed-effects models, using the *lmerTest* R package (16). Models were fitted using maximum likelihood estimation. Cell donor age group (young or old), testosterone treatment and the age x treatment interaction were included as fixed effects. Cell donor and well number nested within cell donor were treated as random effects in all models. Additionally, image number nested within well and cell donor was treated as a random effect for measures of nuclear morphology, whole-myoblast AR intensity and AR nuclear localisation. Estimated marginal means and pairwise contrasts were calculated using the *emmeans* R package (17). AR protein content in proliferating myoblasts was analysed using linear mixed-effects models. Protein abundance for each sample was normalised to the pooled reference sample and log_2_-transformed. Cell donor age group, treatment and the age x treatment interaction were included as fixed effects. Cell donor was treated as a random effect. Relative telomere length in proliferating myoblasts was natural-log-transformed and analysed using linear regression. Cell donor age group was included as the predictor, using the young group as the reference group. Model coefficients and estimated marginal mean contrasts are reported with 95% confidence intervals and p values. Significance was set at *p* < 0.05. Results were visualised as box-and-whisker plots and paired donor-level plots using the *ggplot2* R package (10). For AR protein content, model estimates and 95% confidence intervals were back-transformed to fold changes for interpretation.
